# Widespread transcriptional scanning in the testis modulates gene evolution rates

**DOI:** 10.1101/282129

**Authors:** Bo Xia, Yun Yan, Maayan Baron, Florian Wagner, Dalia Barkley, Marta Chiodin, Sang Y. Kim, David L. Keefe, Joseph P. Alukal, Jef D. Boeke, Itai Yanai

## Abstract

The testis expresses the largest number of genes of any mammalian organ, a finding that has long puzzled molecular biologists. Analyzing our single cell transcriptomic maps of human and mouse spermatogenesis, we provide evidence that this widespread transcription serves to maintain DNA sequence integrity in the male germline by correcting DNA damage through “transcriptional scanning”. Supporting this model, we find that genes expressed during spermatogenesis display lower mutation rates on the transcribed strand and have low diversity in the population. Moreover, this effect is fine-tuned by the level of gene expression during spermatogenesis. The unexpressed genes, which in our model do not benefit from transcriptional scanning, diverge faster over evolutionary time-scales and are enriched for sensory and immune-defense functions. Collectively, we propose that transcriptional scanning modulates germline mutation rates in a gene-specific manner, maintaining DNA sequence integrity for the bulk of genes but allowing for fast evolution in a specific subset.

## Main texts

The testis has been known for many years as the organ with the most complex transcriptome^1–4^. Widespread transcription in the testis has been reported to cover over 80% of all protein-coding genes in human as well as in other species^3–6^. Several hypotheses have been put forth to explain this observation^7,8^. Widespread expression may represent a functional requirement for the gene-products in question^2,7^. However, more complex organs – such as the brain – do not exhibit a corresponding number of expressed genes, despite their significantly greater number of cell types^3–5,9^. Moreover, recent studies have shown that many testis-enriched and evolutionarily-conserved genes are not required for male fertility in mice^10^. The notable discordance between the transcriptome and the proteome in the testis^11,12^ further supports the notion that the widespread transcription does not exclusively lead to protein production via the central dogma.

A second hypothesis implicates leaky transcription during the massive chromatin remodeling that occurs throughout spermatogenesis^7,13,14^. However, this model predicts more expression during later stages of spermatogenesis – when the genome is undergoing the most chromatin changes – in contradiction with previous observations^5,13,15^. Additionally, one would expect leaky transcription to be under tighter control given the high energetic requirements of widespread transcription^16–18^.

Here we propose the ‘transcriptional scanning’ model, whereby widespread testis transcription modulates gene evolution rates. Using scRNA-Seq of human and mouse testes, we confirmed that widespread transcription indeed originates from the germ cells as opposed to a mixture of somatic and germline expression. We next found that spermatogenesis-expressed genes have fewer germline variants in the population compared to the unexpressed genes, and that the signature of transcription-coupled repair (TCR) on these genes could explain the observed pattern of biased germline mutations. Our model of transcriptional scanning suggests that widespread transcription during spermatogenesis acts as a DNA scanning mechanism that systematically detects and repairs bulky DNA damage through TCR^19–21^, thus reducing germline mutations rates and, ultimately, the rates of gene evolution. Genes unexpressed in the male germline do not constitute a random set. Rather, they are enriched in sensory and immune/defense system genes, consistent with previous observations that these genes evolve faster^22–24^. However, transcription-coupled damage (TCD) overwhelms the effects of TCR in the small subset of very highly expressed genes, which are enriched in spermatogenesis-related functions, implicating also a role for TCD in the modulation of germline mutation rates^25^. Collectively, our ‘transcriptional scanning’ model exposes a hitherto unappreciated aspect of DNA repair in biasing gene evolution rates throughout the genome.

### Single-cell RNA-Seq reveals the developmental trajectory of spermatogenesis

To identify the precise gene expression patterns across spermatogenesis we applied single-cell RNA-Seq to the human and mouse testes (Supplementary Fig. 1a)^27^. The resulting data allowed us to distinguish between the genes expressed in the somatic and germline cells, as well as to reveal the dynamic genes expressed throughout the developmental process of spermatogenesis which includes mitotic amplification, meiotic specification to generate haploid germ cells, and finally, differentiation and morphological transition to mature sperm cells (Fig. 1a-b)^28–30^.

**Fig. 1:**
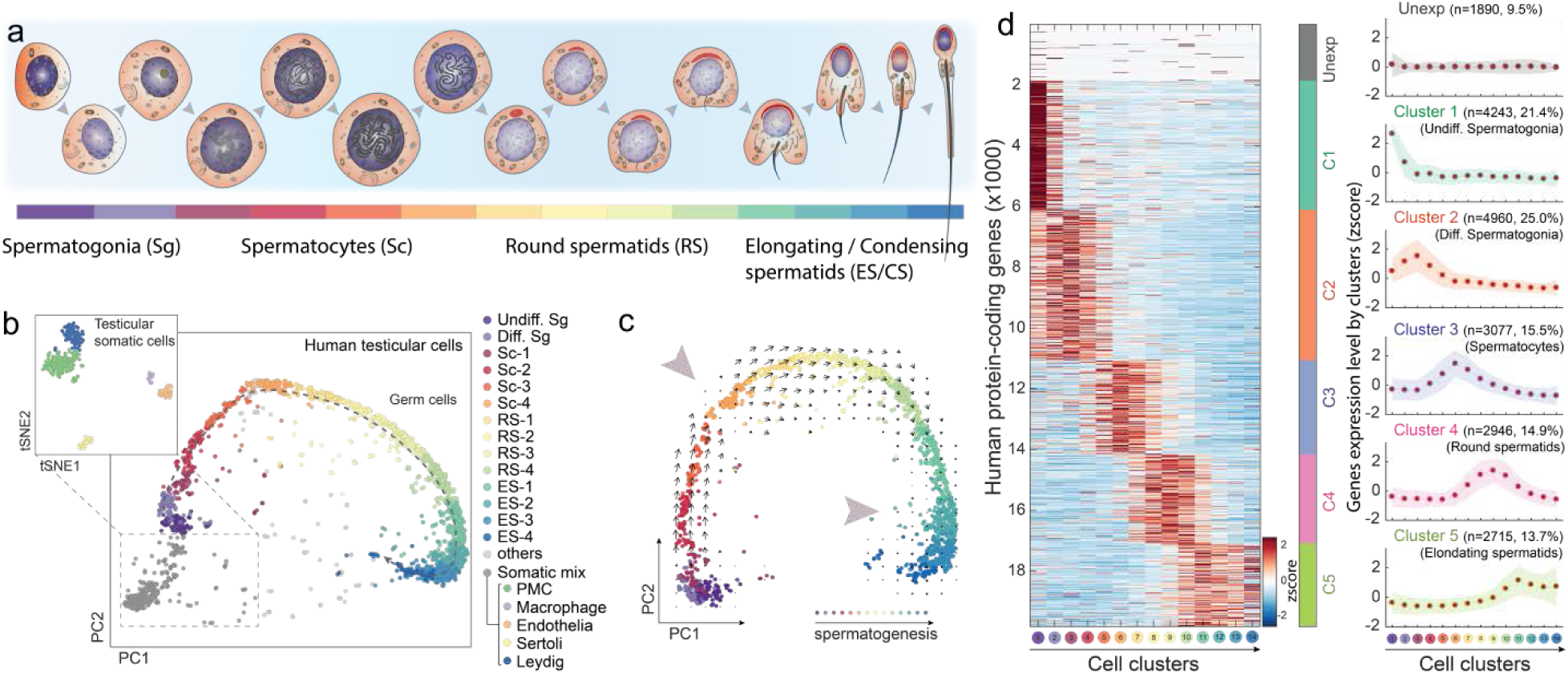
scRNA-Seq reveals a detailed molecular map of human spermatogenesis. **a**, Schematic of developmental stages of human spermatogenesis. **b**, Dimension reduction analysis (PCA and tSNE) of human testes scRNA-Seq results. Colors indicate the main spermatogenic stages and somatic cell types, as defined by unsupervised clustering and marker genes (Supplementary Fig. 1 and Methods). **c**, PCA on the spermatogenic-complement of the single-cell data. Arrows and large arrowheads indicate the RNA velocity algorithm^26^ predicted developmental trajectory and transcriptionally inactive stages during spermatogenesis, respectively (Methods). **d**, Heatmap (left) and plots (right) of the expression patterns of all human protein-coding genes throughout spermatogenesis according to k-means method-defined gene clusters, including the unexpressed gene cluster. The genes numbers and enriched spermatogenesis stage of each cluster are also indicated.

A principal component analysis (PCA) and unsupervised clustering method on the scRNA-Seq data of human testicular cells revealed 19 cells clusters composed of cells from different biological and technical replicates (Fig. 1b and Supplementary Fig. 1b, Methods). We first annotated the 5 cell clusters composed of somatic cells – including Leydig cells, Sertoli cells, peritubular myoid cells, testicular endothelia cells and testis-resident macrophages ^30^ – using previously determined cell type markers (Fig. 1b and Supplementary Fig. 1c-d, Methods). Excluding the somatic cells, PCA on the 14 clusters of germ cells revealed a continuous spectrum suggesting that the order of the cells corresponds to the developmental trajectory of spermatogenesis (Fig. 1c). Three independent lines of evidence support this inference. First, the order of expression of known marker genes across the continuous cluster matches their developmental order (Supplementary Fig. 1d). Second, pseudotime analysis using Monocle2 revealed the same cell trajectory (Supplementary Fig. 1d-e)^31^. Finally, RNA Velocity analysis^26^ – examining the relationship between the spliced and unspliced transcriptomes – further supported the developmental progression during spermatogenesis and also identified the previously reported slowdown of expression during meiosis and late spermiogenesis (Fig. 1c)^13,30^. We thus concluded that germ cell transcriptomes could be ordered as successive stages throughout spermatogenesis.

Our scRNA-seq results on testicular cells allowed us to test whether the long-observed widespread gene expression in the testis has contributions from both germ and somatic cells, or the expression is mainly from the germ cells. Examining only germ cells, we found that 90.5% of all protein-coding genes are expressed (Methods). In contrast, all the detected somatic cell types collectively express 62.3% of the genes. The detailed delineation of spermatogenic trajectory provides stage-specific gene expression with unprecedented resolution (Fig. 1d). To further ask whether specific developmental stages are enriched for expression, we clustered all human protein-coding genes into 6 categories including the unexpressed genes (Fig. 1d, left). The expressed gene sets reflected their enriched expression patterns across all spermatogenesis stages (Fig. 1d, right). While no single stage accounts for the widespread transcription, we can infer that each cell will gradually express the observed ∼90.5% of the genes throughout its overall maturation to a sperm.

To test the generality of these results, we repeated the experiments on mouse testis samples and found that the pattern of transcription during mouse spermatogenesis was broadly comparable to that of human (Supplementary Fig. 2a-d). In terms of genes expressed across the stages, we found an overall highly conserved spermatogenesis gene expression program (Supplementary Fig. 2b-d). A combined principal component analysis of human and mouse germ cells further highlighted this conserved transcriptional program of spermatogenesis (Supplementary Fig. 2e-g). Interestingly, PC2 clearly separates the human and mouse cells (Supplementary Fig. 2h), indicating a species-specific gene expression signature between the two species. These genes include metabolic genes like *GAPDH* (*Gapdh*)^32^ and *FABP9* (*Fabp9*)^33^, chemokine gene *CXCL16* (*Cxcl16*), and sperm motility-related gene *SORD* (*Sord*)^34^ (Supplementary Fig. 2i). Collectively, these results highlight the overall gene expression conservation of human and mouse spermatogenesis, but also identified the divergence between the two species.

### Reduction of germline mutation rates in spermatogenesis expressed genes

We hypothesized that widespread transcription during spermatogenesis could lead to two scenarios (Fig. 2a): 1) transcription events unwind the double-strand DNA, leading to an increased likelihood of mutations by transcription-coupled damage (TCD)^25^, and consequently to higher germline mutation rates and diversity within the population; and/or 2) the transcribed regions are subject to transcription-coupled repair (TCR) of DNA damages^19–21^, thus reducing germline mutation rates and safeguarding the germline genome, leading to lower population diversity. In both scenarios, differences in expression states may contribute to the pattern of population diversity, and ultimately lead to differential gene evolution rates.

**Fig. 2:**
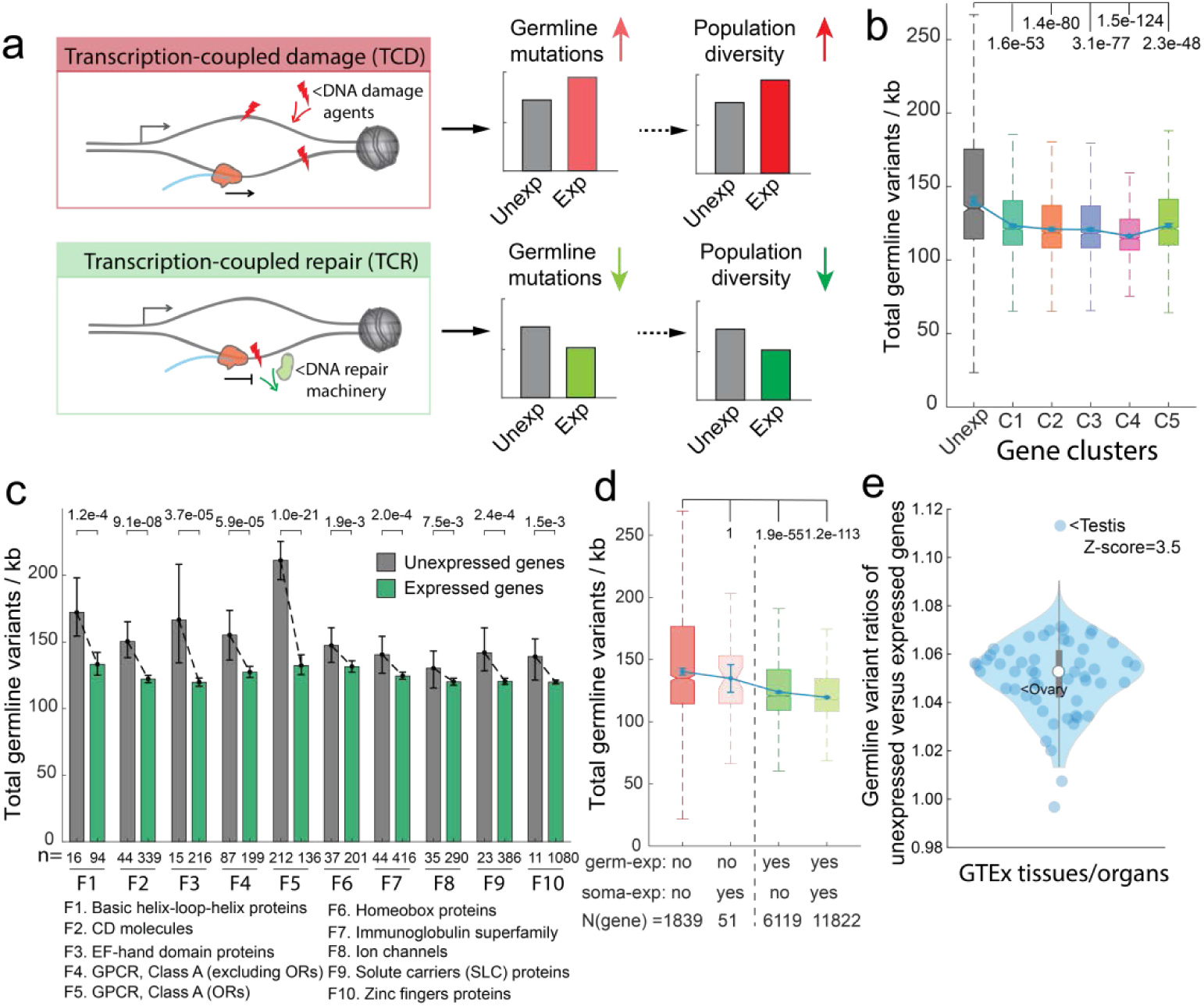
Widespread transcription in spermatogenic cells is associated with reduced germline mutation rates. **a**, Two possible consequences of widespread transcription in spermatogenic cells. **b**, Total germline variant levels across the gene clusters, as determined in Figure 1d. **c**, Total germline variant levels of expressed and unexpressed genes across large gene families (Methods). **d**, Total germline variant levels across gene sets as determined by binarized expression (expressed versus unexpressed) in testicular germ cells and the somatic cells. **e**, Ratios of germline variants in unexpressed and expressed genes in diverse human organs and cell types. Dot represents individual tissues/organs from the GTEx-project^35^. Significance in (**b**-**d**) is computed by the Mann-Whitney test between expressed and unexpressed gene sets with Bonferroni correction for multiple tests. Error bars indicate 99% confidence intervals calculated by bootstrap methods with n=10,000 (Methods).

The public databases have amassed over 200 million germline variants detected in the human population, providing a rich resource for studying germline mutation rates^36^. Since ∼80% of these germline variants are thought to have originated in males^37,38^, we used this dataset to query for the predicted effects caused by widespread transcription, according to the two scenarios^39–41^. Interestingly, we found that spermatogenesis-expressed genes, regardless of the timing of their expression (throughout and following meiosis), generally have a lower level of germline mutations, relative to the unexpressed genes (Fig. 2b), consistent with the previous notion of transcription-coupled repair in spermatogenic cells ^42,43^. This difference is robust across variations in gene clustering and individuals (Supplementary Fig. 3a-e) and is not observed in the gene flanking sequences (5kb upstream and downstream), indicating a genic region-specific effect (Supplementary Fig. 3f-h).

To further control for differences in DNA variation specific to particular sequence domains of genes expressed in the germ cells, we examined gene families individually according to germline expressed (in any stage) and unexpressed groups (Methods)^44^. For all large gene families (>100 genes) with at least 10 genes in either category we found lower germline variants level in the spermatogenesis-expressed gene subgroup (Fig. 2c). For example, of the 110 genes with a basic helix-loop-helix domain, 94 are expressed in the germ cells, and the expressed subgroup has a 22% reduction in germline variant level in the population as compared to the unexpressed complement.

We further controlled for the uniqueness of this effect to male germline gene expression, relative to that of other cell types. By distinguishing the binarized expression status in both germ cells and testicular somatic cell types (Fig. 1b), we found that genes expressed exclusively in somatic cells do not exhibit a reduced germline mutations (Fig. 2d, Methods). To study somatic tissues more broadly we turned to the Genotype-Tissue Expression (GTEx) dataset which has characterized transcriptional profiles across all major human tissues/organs, including testis^35^. While not at the single-cell level and thus effectively averaging across cell types, testis expression in this dataset showed a significant difference relative to all other tissues in its reduction between the expressed and unexpressed gene complement (Z-score = 3.5; Fig. 2e). Interestingly, we found that the ovary transcriptome does not predict such an effect, consistent with the notion that point mutations mainly originate in male germ cells^37,38^. Altogether, these results support the second scenario of transcription-coupled repair in the male germ cells (Fig. 2a), with male germ cell-expressed genes showing reduced levels of germline mutations rates.

### The signature of TCR of the germline mutations of spermatogenesis-expressed genes

If the reduction of mutations follows from a TCR-induced process, we would expect an asymmetry between the germline mutation levels of the coding and the template strands in the spermatogenesis expressed genes^43,45–48^, but not in the unexpressed genes (Fig. 3a). The asymmetry would be such that the template strand accumulates fewer mutations since, in TCR, DNA damage is detected by the RNA polymerase on the template strand^19^. To distinguish between mutations occurring on the coding and template strands, we adapted previous approaches to identify strand-asymmetries in the mutation rate (Fig. 3b)^43,45^. By studying mutation categories with reference to the coding and template strand, Haradhvala *et al.* inferred a bias in somatic mutation rates^45^ and such a strategy was also utilized by Chen *et al* ^43^. We applied this approach to germline mutations and found that a lower mutation rate was inferred to occur on the template strands of expressed genes during spermatogenesis, regardless of its expression pattern along the spermatogenesis stages, while such an effect was not apparent in the unexpressed genes, as represented by A-to-T (A>T) transversion mutations in Figure 3c and in the other mutation types (Supplementary Fig. 4b). Interestingly, we found that the coding strand of the expressed genes also has a lower mutation rate than the coding strand of the unexpressed genes (Fig. 3c and Supplementary Fig. 4b), suggesting that antisense transcription in spermatogenesis might further reduce mutation levels^49^.

**Fig. 3:**
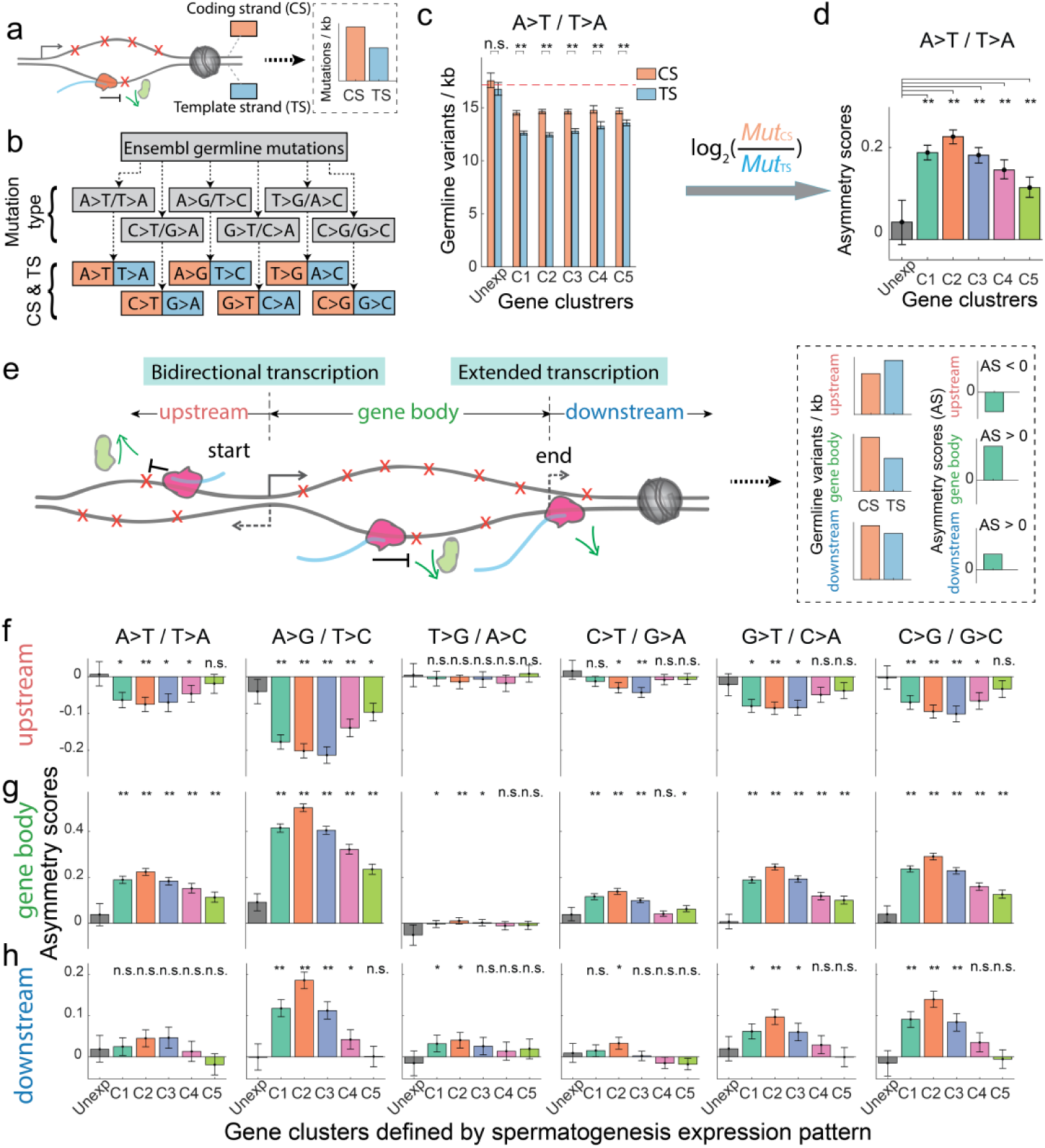
TCR-associated mutation asymmetry scores show bidirectional transcription and extended transcription signatures. **a**, Schematic of a transcribed gene with the template strand containing lower DNA damage and, consequently, a lower mutation rate. **b**, Germline mutations associated with genes were retrieved from Ensembl^36^, classified into the six mutation classes, and further distinguished in terms of coding and template strands, as previously introduced^45^. **c**, A-to-T transversion mutation rates for the coding and the template strands for the spermatogenic gene categories. Dashed lines indicate the average level of mutations in the unexpressed genes. **d**, Asymmetry scores throughout spermatogenic gene categories, computed as the log2 ratio of the coding to the template mutation rates (shown in **c**). **e**, Schematic of gene architecture indicating bidirectional and extended transcription. The schematic shows that relative to the promoter, upstream and gene body transcription occur on opposite strands, while downstream transcription occurs on the same strand as the gene body. **f-h**, Asymmetry scores in the upstream 5kb region (**f**), gene body (**g**) and downstream 5kb region (**h**) across all six mutation types. Significance between the unexpressed gene category and the expressed gene categories (**d**, **f**-**h**) and between coding and template strands (**c**) was computed by the Mann-Whitney test with Bonferroni correction. *, *P*<0.01; **, *P*<0.000001; n.s., not significant. Error bars indicate 99% confidence intervals calculated by bootstrap methods with n=10,000.

We next computed an ‘asymmetry score’ to study the ratio between mutation levels inferred to occur in the coding and template strands (Fig. 3c-d)^45^. As expected, the unexpressed group of genes has minimal asymmetry score levels (Fig. 3d and 3g), indicating an absence of transcription-induced removal of DNA damage. As negative controls, we found that mutational asymmetry was not observed when comparing Watson and Crick strands (instead of gene-specific coding and template strands, Supplementary Fig. 6), nor did we detect difference between the gene clusters when shuffling the spermatogenic gene clustering assignments (while maintaining the group sizes, Supplementary Fig. 7).

### Bidirectional transcription signatures of mutation asymmetries

While our analysis thus far examined transcription in the gene body (i.e. genic region of transcription start site to the end site, also referred as the transcription unit), transcription in the human genome contains additional levels of complexity. In particular, though transcription is usually considered in the gene body, initiation can occur on the opposite strand, leading to upstream transcription in the opposite direction^50,51^ (Fig. 3e). If lower mutation rates are indeed transcription-induced, we predicted that mutation asymmetry scores would display an inverse pattern between the opposite sides of the initiation of bidirectional transcription (Fig. 3e). Consistently, we detected an inverse pattern of asymmetry scores between the gene body and the upstream sequences (Figs. 3f-g, Supplementary Fig. 4a-b). Furthermore, since transcription may extend beyond the annotated end or alternative polyadenylation sites (Fig. 3e)^52^, we also predicted that the asymmetry scores in the downstream sequences would display a similar, though expectedly weaker pattern compared to that of the gene body (Fig. 3e). Again, we found the expected pattern in which the gene body and the downstream sequences have the same pattern of asymmetry scores (Figs. 3g-h, Supplementary Fig. 4b-c).

Finally, we predicted that the same TCR influences would be manifested in the mouse data, and indeed found such evidence (Supplementary Fig. 8). For example, G-to-T (G>T) transversion mutations show strong conserved asymmetric mutation patterns in both the human and mouse data. Since G-to-T mutations come predominantly from endogenous oxidative DNA damage of guanine^40,53^, such conserved asymmetric germline mutation patterns between gene coding and template strands further support the notion of TCR effects on germline mutations. Collectively, these analyses provide support for transcription-induced germline mutation reduction in spermatogenesis expressed genes.

### Transcriptional scanning is tuned by gene-expression level

Our results led us to propose ‘transcriptional scanning’ as a mechanism to systematically reduce DNA damage-induced mutagenesis in the bulk of genes by widespread spermatogenic transcription to safeguard the germline genome sequence integrity (Fig. 4a). Such a mechanism suggests that mutation rates of scanned genes might be tuned by their expression levels in the testis. To test this, we binned all genes into nine groups according to their expression levels (Fig. 4b, Methods). Consistently, we found that even the most lowly-expressed genes have lower levels of germline mutations than the unexpressed genes (Fig. 4c-d).

**Fig. 4:**
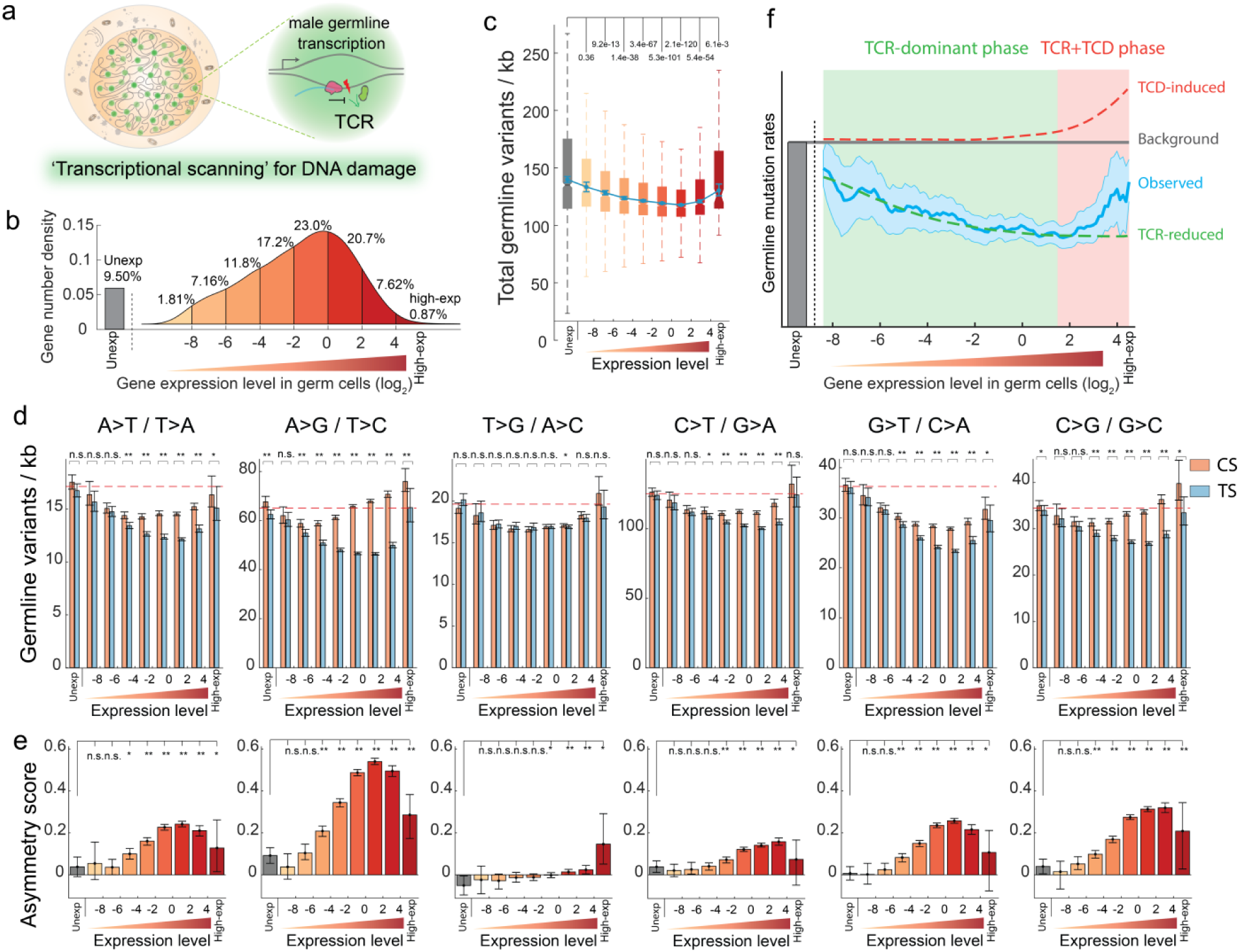
‘Transcriptional scanning’-induced mutation reduction is tuned by gene-expression levels. **a**, Schematic of transcriptional scanning of DNA damage in male germ cells. **b**, Genes were binned to nine expression level groups, from unexpressed (Unexp) to highly expressed (High-exp) (Methods). **c**, Germline mutation rates across gene expression level categories. Spermatogenesis unexpressed- or highly expressed-genes have higher level of germline mutations. **d**, Distributions of the indicated germline mutation types across gene expression level categories, and distinguished by coding and template strands. Dashed lines indicate the average level of mutations in the unexpressed genes. **e**, Distribution of asymmetry scores between coding and template strand for the mutation types indicated in **(d)**. **f**, Expression level tuning of germline mutation rates following additive contributions by transcription-coupled repair (TCR-reduced) and transcription-coupled damage-induced (TCD-induced) effects. The observed germline mutation level represents average mutation rates across 100 evenly-binned expression levels, with background shadows indicating 99% confidence intervals of expression level-associated germline mutation rates. Significance between the unexpressed gene category and the expressed gene categories (**c** and **e**) or between germline variants on coding strand and template strand (**d**) is computed by the Mann-Whitney test with Bonferroni correction. *, *P*<0.01; **, *P*<0.000001; n.s., not significant. Error bars indicate 99% confidence intervals calculated by bootstrap method with n=10,000.

‘Transcriptional scanning’ predicts that higher expression levels would lead to additional scanning, and, consequently, to further reduced mutation rates on the template strand. Indeed, examining our asymmetry score according to different expression levels, we observed that as expression level increases, the overall mutation level drops (Fig. 4c-d). Surprisingly, however, the very highly expressed genes showed the opposite effect: asymmetry between the strands is reduced and a higher level of germline mutations relative to the moderately expressed genes is observed (Figs. 4c-e). This pattern is consistent, however, with observations that very high expression levels can lead to transcription-coupled DNA damage (Fig. 2a), as previously reported for transcription-associated mutagenesis in highly expressed genes in other systems^25,54^. The mutation type in which TCD is most evident is A-to-G transitions (Fig. 4c), and similarly, such strong TCD-induced effect was readily observed in somatic A-to-G mutations in liver cancer samples^45^. Together, the TCD-induced effect in the very highly expressed genes during spermatogenesis persists across all mutation types (Fig. 4d-e).

Overall, our analyses suggest that spermatogenesis gene expression levels tune germline mutation levels by ‘transcriptional scanning’ which reduces mutation rates in genes with low-expression (Fig. 4f). Increasing expression levels are correlated with further reductions in mutation rates, but only to a point. In the very highly expressed genes, TCD overwhelms the TCR-induced reductions, and produces an overall higher germline mutation rate than genes expressed at low and moderate levels (Fig. 4f).

### The role of transcriptional scanning in genome evolution

Since the ‘transcriptional scanning’ mechanism is proposed to reduce germline mutations, we asked why any gene would be unexpressed during spermatogenesis, instead of benefiting from this process. Studying the set of 1,890 unexpressed genes at the functional level, we observed enrichment for environmental sensing, immune systems, defense responses, and signaling functions (Fig. 5a and Supplementary Table 1). These functions coincide with those known to be fast-evolving in the human genome^22–24^, suggesting that their lack of expression in the testis is related to their evolution. Consistently, we detected the highest rates of sequence divergence across ape genomes in the unexpressed genes (Fig. 5b, Supplementary Fig. 9a). While selection is typically invoked to account for the fast evolution of genes (Supplementary Fig. 9b-c), biased germline mutation rates may also contribute according to the neutral theory of gene evolution^22,23,41,55–57^. To test this, we studied the synonymous substitution rates (dS, generally assumed to be neutral) as a proxy for the germline mutation rates and used this measure to compare between the spermatogenesis expressed and unexpressed genes. Interestingly, we found that the spermatogenesis-expressed genes have lower dS values, supporting the notion that biased germline mutation rates also contribute to the biased gene evolution rates. We further found that the very highly expressed genes in spermatogenesis have increased rates of divergence (Supplementary Fig. 9f-i). As expected from their high expression, we found that this set of genes is mainly enriched for roles in male reproduction (Supplementary Fig. 9j and Table 2).

**Fig. 5:**
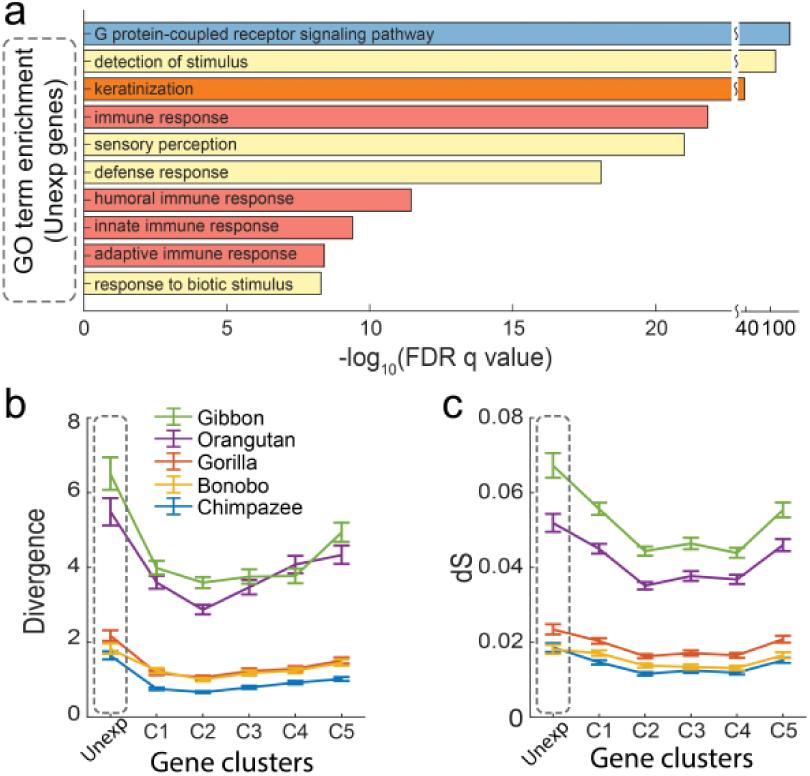
Evolutionary consequences of transcriptional scanning in male germ cells. **a**, Gene ontology terms enriched in the set of genes unexpressed during spermatogenesis. ‘FDR q-value’ indicates the GO term enrichment significance test p-values after multiple-test correction by the Benjamini-Hochberg method^58^. **b-c**, DNA divergence levels **(b)** and dS scores **(c)** and of human genes with their orthologous in the indicated apes, according to gene expression-pattern clusters. Gray dashed box highlights the male germ cell-unexpressed gene cluster.

To disentangle the effects of DNA repair and selection on the different gene evolution rates between spermatogenesis expressed and unexpressed genes, we compared the frequency of variants across introns and coding sequences (CDS), where we expect most variants to be neutral and a mix of neutral and under selection, respectively. Consistent with the dS results, we found an average reduction of 5.49% when comparing the intron variant levels between the expressed and the unexpressed genes. In contrast, for the CDS region variants, we observed a reduction of 9.56%, likely reflecting the mixed effects of transcriptional scanning and natural selection (Supplementary Fig. 9d-e). These data suggest that the unexpressed genes are under a unique selection regime whereby occurring mutations are less likely to be purged by purifying selection. Together, our results suggest that, beyond selection, transcriptional scanning in spermatogenesis imposes an additional bias in modulating gene evolution rates.

## Discussion

Our findings led us to propose the ‘transcriptional scanning’ model, whereby widespread transcription in spermatogenesis leads to a rugged landscape of biased germline mutations (Fig. 6a). In this model, widespread transcription in the testis acts to systematically reduce germline mutations by transcription-coupled repair (TCR), thereby safeguarding the germ cell genome sequence integrity. Given that this process is carried out in the germline, the variable mutation rates have important implications. Over evolutionary time-scales, combined with natural selection, the spermatogenesis-expressed genes evolve more slowly (Fig. 6a, middle). The small group of genes that are unexpressed in spermatogenesis are enriched for sensory and immune/defense system genes (Fig. 5c) and exhibit higher mutation rates, which is explained in our model by their lack of TCR-mediated germline mutation reduction (Fig. 6a, left). Immune and defense system genes are known to evolve faster^22–24^ and our biased transcriptional scanning model provides insight into how variation is preferentially provided to this class of genes. Such biased germline mutation rates provide increased population-wide genetic diversity which may be under strong selective biases for adaptation at the population-level in rapidly changing environments. A third class of genes with very high germline expression exhibit higher germline mutation rates since their transcription-coupled DNA damage obscures the effect of TCR (Fig. 4f and Fig. 6a right). This model provides a more comprehensive view of the combined effects of TCR and TCD in spermatogenic cells (Fig. 4f), and refines previous observations that germline mutation rates increase with expression levels while highly expressed genes evolve slower^43,54,59,60^. While the observed mutational bias does not alone direct evolution according to our model – since fixation in the population is also influenced by genetic drift and natural selection – it is expected to contribute to global gene evolution rates.

**Fig. 6:**
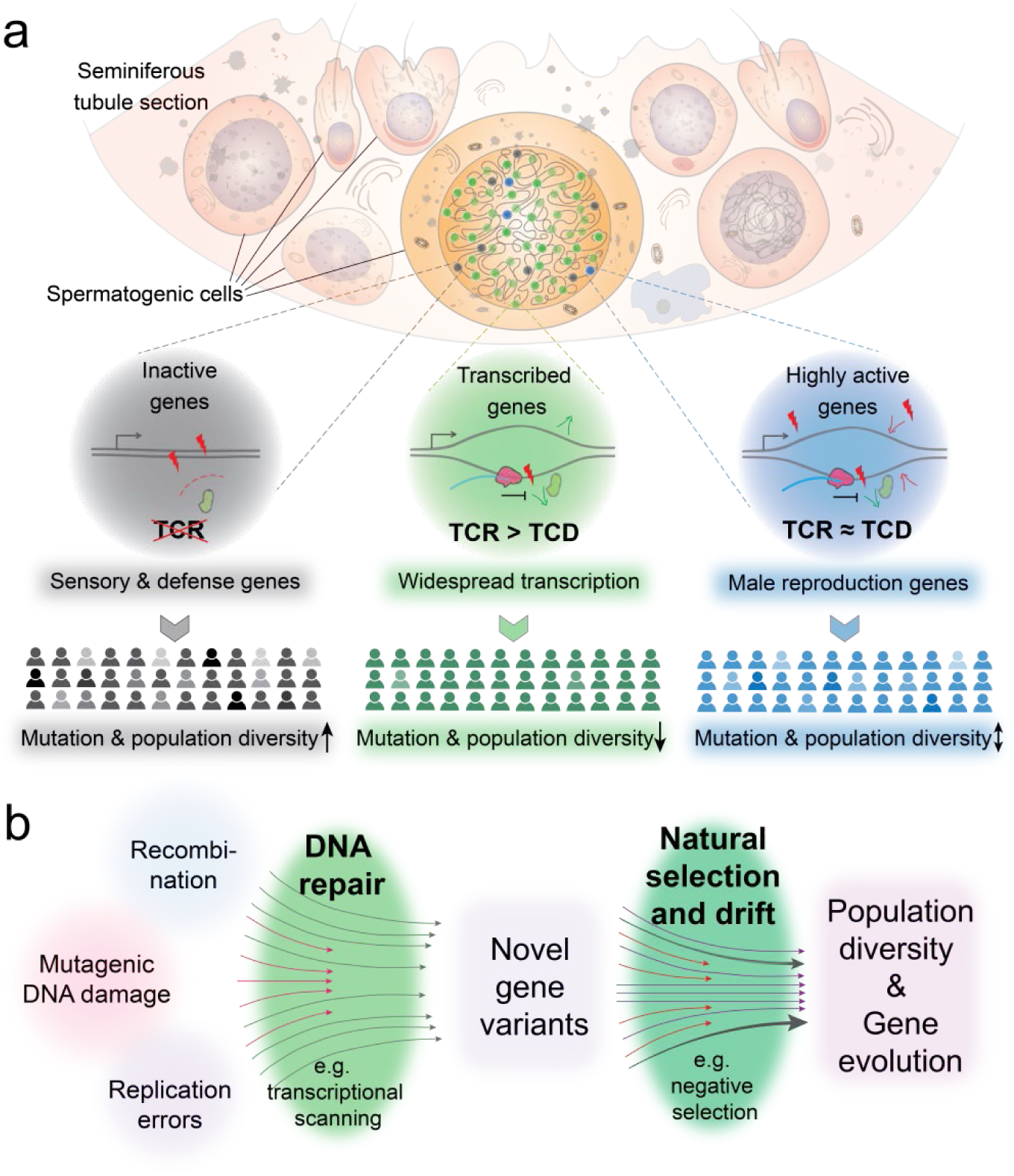
A model for widespread transcriptional scanning in male germ cells and its contribution to gene evolution. **a**, The transcriptional scanning model predicts reduced germline mutation rates across most expressed genes. Genes unexpressed in spermatogenesis have relatively higher mutation rates and consequently experience more evolutionary divergence. In the very highly-expressed genes, transcription-coupled DNA damage overwhelms the effects of TCR, resulting in higher mutation rates in these genes, highly enriched for male reproductive function genes. **b**, A revised model for generating DNA sequence variation and gene evolution. DNA repair, represented by transcriptional scanning, acts as a biased mechanism for the generation of novel DNA variants and, ultimately, to gene evolution rates.

Gene sequence evolution requires (1) the generation of novel DNA variants, stemming from DNA damage-induced mutagenesis, replication errors and/or recombination, and (2) natural selection and/or drift on the novel variants^41,57^. Since our results implicate a DNA-repair mechanism in biasing the production of variants throughout the genome, we propose that this represents a hitherto unappreciated aspect in the establishment of differential gene evolution rates. Thus, DNA repair pathways act to constrain mutagenic DNA damage in a biased manner, analogous to the effects of selection and drift in the population (Fig. 6b). By understanding these patterns of uneven germline mutations and the intrinsic removal mechanism of germline DNA damage, our model provides insight into mutation-driven genome evolution^61^.

While transcriptional scanning is proposed to systematically detect and remove bulky germline DNA damage, male germ cells are still expected to retain mutations that cannot be repaired by the TCR machinery^20,62,63^. These male germline mutations likely originate from DNA replication errors, accumulating with paternal age^64^, or less bulky DNA damages like base deamination^65^. Thus, it will be of interest to analyze germline mutation pattern with a focus on other signature mutation types beyond TCR45,65,66.

Our model leads to important testable predictions and may provide deeper insights into human genetics and diseases. First, our model predicts that male-derived *de novo* mutations should occur more frequently in genes that are unexpressed during spermatogenesis. Second, the same process should also hold in other species which have readily observed similar widespread transcription in male germ cells^5^, as we also provide evidence for conserved transcriptional scanning in mouse (Supplementary Fig. 8). Finally, we expect TCR-deficient animals to produce offspring with an increase in the number of *de novo* mutations and that they should not show that characteristic lower mutation rates in the template – versus the coding – strand. For patients with TCR gene-associated mutations, such as Cockayne syndrome and xeroderma pigmentosum^67^, our model predicts overall higher germline mutation rates. Lastly, embryonic stem cells (ESCs) share similar patterns of widespread transcription^68^, leading us to speculate that systematic scanning and removal of DNA damage also functions in ESCs. If so, beyond spermatogenesis, transcriptional scanning may be deployed to achieve lower mutation rates in ESCs and in the early developing embryos^68–70^.

## Acknowledgments

We thank Yael Kramer, Xavier Sanchez and Fang Wang for coordinating the human testicular tissue collection. We thank Molly Przeworski, Hannah Klein, Huiyuan Zhang and the members of the Yanai lab for constructive comments and suggestions to the manuscript. We thank Megan Hogan, Raven Luther and Matthew Maurano for assistance with sequencing. This work was supported by the NYU School of Medicine with funding to I.Y.

## Author contributions

B.X. and I.Y. conceived the project, interpreted the results and drafted the manuscript. B.X. led the experimental and analysis components. Y.Y. contributed to the RNA velocity and pseudotime analysis, and processed the raw germline variants data from the public database. M.B., D.B. and M.C. contributed expertise in the inDrop experiments. M.B. and F.W. helped with scRNA-seq data analysis. F.W. built the inDrop sequencing data mapping pipeline. J.A., S.Y.K., and D.K. contributed to the sample collection. J.B. contributed to interpreting the results. All authors edited the manuscript.

## Methods

### Human testicular tissue

Human testicular tissue was obtained from New York University Langone Health (NYULH) Fertility Center; this was approved by the NYULH Institutional Review Board (IRB). Fresh seminiferous tubules were collected separately from testicular sperm extraction (TESE) surgery of two healthy patients with an obstructive etiology for infertility; there were no drug or hormonal treatments prior to TESE surgery. The research donors were fully informed before signing consent to donating excess tissue for research use; this was again done in fashion consistent with the IRB (including tissue sample de-identification).

### Human testicular single cell suspension preparation

After TESE surgery, samples were kept in cell culture PBS and transported to the research lab on ice within 1h of surgery for single-cell preparation. Testicular single-cell suspension was prepared by adapting existing protocol^71^. Specifically, samples from TESE surgery was washed once with PBS and resuspended in 5mL PBS. Seminiferous tubules were minced quickly in a cell culture dish and spun down at 100g for 0.5min to remove supernatants. The minced tissue was resuspended in 8mL of 37°C prewarmed tissue dissociation enzyme mix (See below). Tissue dissociation was done by incubating at 37°C for 20min with mechanical dissociation with pipetter every 5min. After digestion, the reaction was quenched by adding 2mL of 100% FBS (Gibco, Cat. 16000044) to a final concentration of 10%. Dissociation mix was filtered through a 100um strainer to remove remaining seminiferous tubule chunks. Cells were washed once with DMEM medium (Gibco, Cat. 11965092) with 10% of FBS and twice with PBS to remove residual EDTA. Cell viability was checked with Trypan-blue staining (with expectation of over 85% viable cells) before moving to the inDrop microfluidics platform. The tissue dissociation enzyme mix (8mL) was composed of 7.56mL of 0.25% Trypsin-EDTA (Gibco, Cat. 25200056), 400uL of 20mg/mL type IV Collagenase (Gibco, Cat. 17104019) and 40uL of 2U/uL TURBO DNase (Invitrogen, Cat. AM2238).

### Mouse testicular single cell preparation

C57BL/6J mice (4-month old) were bought from the Jackson Laboratory through the New York University Langone Health (NYULH) Rodent Genetic Engineering Laboratory. Mice were anesthetized before sacrificing for testicular tissue collection following the NYULH IRB requirements for experimental animal operation. The dissociated testicular tissue was kept in the PBS buffer and then transported to the research lab on ice immediately for single-cell dissociation. The tissue dissociation protocol is slightly different from the human testicular tissue dissociation. The whole testis was decapsulated in PBS buffer to collect the seminiferous tubules. The seminiferous tubules were quickly minced into small pieces of ∼2-5mm and then washed once with PBS buffer. The minced tissue was resuspended in 8mL of 37°C pre-warmed tissue dissociation buffer 1 (1mg/mL type IV Collagenase in DMEM medium) and incubate at 37°C for 5min. This pre-dissociation step removes majority of the interstitial cells. The tissue was then spun down at 100g for 1min to remove supernatants. The tissue was resuspended by 8mL tissue dissociation buffer 2 (7.96mL of 0.25% Trypsin-EDTA and 40uL of 2U/uL TURBO DNase). The second tissue dissociation was done by incubating at 37°C for 15min with mechanical dissociation with pipetter every 5min. The dissociation was quenched by adding 2mL of 100% FBS to a final concentration of 10%. Dissociation mix was filtered through a 100um strainer to remove any remaining tissue chunks. Cells were washed once with DMEM medium and twice with PBS to remove residual EDTA. Cell viability was checked with Trypan-blue staining (both replicates have over 95% viable cells) before moving to the inDrop microfluidics platform.

### Single-cell RNA-Seq

Single-cell barcoding was carried out with the inDrop microfluidics platform^27^ as instructed by the manufacturer (1CellBio). Briefly, the microfluidic chip and barcoded hydrogel beads were primed ahead of single cell preparation. The ready-to-use single-cell suspension in PBS (after two times wash with PBS buffer) was adjusted to 0.1 million/mL by counting with hemocytometer. Next, the prepared cells, reverse transcription reagents (SuperScript III Reverse Transcriptase, Invitrogen, Cat. 18080085), barcoded hydrogel beads and droplet-making oil were loaded onto the microfluidic chip sequentially. Encapsulation was done by adjusting microfluidic flow rates as instructed. Single-cell barcoding and reverse transcription in the droplets were done by incubating at 50°C for 2h followed by heat inactivation at 70°C for 15min. Then the droplets containing barcoded single-cells were aliquoted aiming for 1000-2000 cells per aliquot and then decapsulated by adding demulsifying agent.

### Sequencing library preparation

Single-cell RNA-Seq library preparation after inDrop was carried out as instructed by the manufacturer (1CellBio) and similar to the CEL-Seq2 method^72^. Basically, barcoded single-cell cDNA was purified with Agencourt RNAClean XP magnetic beads (Beckman Coulter, Cat. A63987) followed by second-strand synthesis reaction with NEBNext mRNA Second Strand Synthesis KIT (New England Biolabs, Cat. E6111S). Then linear amplification of cDNA was carried out through *in vitro* transcription (IVT) using HiScribe T7 High Yield RNA Synthesis kit (New England Biolabs, Cat. E2040S). IVT-amplified RNA was fragmented and purified again with Agencourt RNAClean XP magnetic beads. The second reverse transcription was done with PrimeScriptTM Reverse Transcriptase (Takara Clonetech, Cat. 2680A) followed with cDNA purification with Agencourt AMPure XP magnetic beads (Beckman Coulter, Cat.A63881). cDNA quantity was determined by qPCR on a fraction (5%) of purified cDNA. Final PCR amplification was done according to qPCR results and purified with Agencourt AMPure XP magnetic beads. Library concentration was determined by Qubit dsDNA HS Assay Kit (Invitrogen, Cat. Q32851). Library size was determined by Bioanalyzer High Sensitivity DNA Kit (Agilent, Cat. 5067-4626).

### High-throughput sequencing

Single-cell RNA-Seq library sequencing was carried out with Illumina NextSeq 500/550 75 cycles High Output v2 kit (Cat. FC-404-2005). Custom sequencing primers were used as instructed by manufacturer^27^. In addition, 5% of PhiX Control v3 (Illumina, Cat. FC-110-3001) library was added to give more complexity to scRNA-Seq libraries. Pair-end sequencing was carried out with read1 (barcodes) for 35bp, index read for 6bp and read2 (transcripts) for 50bp.

### Sequencing data processing

Raw sequencing data obtained from the inDrop method were processed using a custom-built pipeline, available at (https://github.com/flo-compbio/singlecell). Briefly, the “W1” adapter sequence of the inDrop RT primer was located in the barcode read (the second read of each fragment), by comparing the 22-mer sequences starting at positions 9-12 of the read with the known W1 sequence (“GAGTGATTGCTTGTGACGCCTT”), allowing at most two mismatches. Reads for which the W1 sequence could not be located in this way were discarded. The start position of the W1 sequence was then used to infer the length of the first part of the inDrop cell barcode in each read, which can range from 8-11 bp, as well as the start position of the second part of the inDrop cell barcode, which always consists of 8 bp. Cell barcode sequences were mapped to the known list of 384 barcode sequences for each read, allowing at most one mismatch. The resulting barcode combination was used to identify the cell from which the fragment originated. Finally, the UMI sequence was extracted, and reads with low-confidence base calls for the sex bases comprising the UMI sequence (minimum PHRED score less than 20) were discarded. The reads containing the mRNA sequence (the first read of each fragment) were mapped to the references genomes (here human GRCh38 and mouse GRCm38) by STAR 2.5.3a with parameter ‘— outSAMmultNmax 1’ and default settings otherwise^73^. Mapped reads were split according to their cell barcode and assigned to genes by testing for overlap with exons of protein-coding genes and long non-coding RNA genes, based on genome annotations from Ensembl release 90. For each gene, the number of unique UMIs across all reads assigned to that gene was determined (UMI filtering), corresponding to the number of transcripts expressed and captured.

### Quality filtering of the scRNA-seq data

Single cells with a total transcript count of less than 1,000 or more than 20% of transcripts originating from either mitochondrial genes (i.e., genes that are part of the mitochondrial genome) or ribosomal protein genes were removed for downstream analysis. After filtering, the single cells from different biological or technical replicate were merged together for downstream analysis. In total, we have 2554 cell from human, with 6499 UMI counts and 2495 detected genes on average. From mouse testis, we obtained 1593 cells in total, with 8998 UMI counts and 2601 detected genes on average.

### Testicular cell clustering and cell type identification

Following quality cell filtering, clustering was done by k-means on the principal component analysis scores, with k determined by ‘elbow-method’^74^. To increase the resolution of cell clustering, the raw UMI counts of testicular single cells were pre-processed through the kNN-smoothing method, with k=3 which indicates a smoothing with the nearest 3 single cell transcriptomes which greatly reduce the noise in scRNA-seq data while retaining the variance between single cells^75^. The principal component analysis used for cell clustering was performed on the smoothed UMI expression matrix of all testicular cells. The pre-processed expression matrices were first normalized to 100,000 transcripts per cell to calculate Fano factor (or variance-to-mean ratio, VMR)^76^. Genes with a Fano factor larger than 1.5 folds of the mean values were defined as dynamically expressed genes. In total, 3615 dynamically expressed genes were selected from the human datasets for downstream PCA visualization and cell clustering. PCA was then performed on the normalized and log_2_ transformed expression matrix using the dynamically expressed genes. Cell clustering was done by k-means clustering with elbow-methods determined k. Following first rounds of cell clustering (k=24), several marker genes were used to determine spermatogenic cell types/states versus somatic cells. *DDX4* (also called *VASA*) was used as a pan-germ cell marker to distinguish the spermatogenic cell lineage. *FGFR3* and *DMRT1*^28,77^ were used to determine spermatogonia. *SYCP3* and *TEX101*^6,78^ were used to determine spermatocytes. *ACRV1* and *ACTL7B*^6,78^were used to determine round spermatids. *TNP1, PRM1, PRM2, YBX1, YBX2* and *HILS1*^6,13,79,80^ were used collectively to determine elongating spermatids states. Together, we identified 14 human spermatogenic cell clusters with at least 50 cells in each cluster (min value as 69 cells, corresponding to spermatocyte-1). Seven cell clusters which overlapped with each other were identified as somatic cells (as shown in Fig. 1b). These cells were isolated for visualization through the t-distributed stochastic neighbor embedding (tSNE) algorithm and re-clustered with an additional k-means clustering algorithm (k=5), as shown in Fig. 1b and Supplementary Fig. 1c. In summary, *CYP11A1, CSF1*, and *IGF1* ^78,81,82^ genes were used to identify Leydig cells; *WT1* and *SOX9*^78,83^ were used to identify Sertoli cells; *MYH11* and *ACTA2* were used to identify peritubular myoid cells^84^; *CD68* and *CD163* were used to identify macrophages^85^; *PECAM1* and *VWF* were used to identify endothelia cells^86^. Three small clusters with mixed expression profiles and/or bad quality were labeled as “other” and discarded as potential contaminants. Mouse testicular cells were analyzed in the same process. In brief, 1915 dynamically expressed genes were selected from the mouse datasets for PCA and cell clustering. Cell clustering with k-means algorithm generated 16 clusters (optimum k defined by elbow-method), out of which 13 clusters were kept as mouse spermatogenic cell clusters, and 3 clusters with few cells were discarded for downstream analysis.

### Pseudotime analysis with Monocle2

We used the R package ‘Monocle2’ (version 2.6.1)^31^ to infer pseudotime tracks for both human and mouse spermatogenic cells. The raw UMI counts of the isolated spermatogenic cells were pre-processed through the kNN-smoothing method (k=3) before performing pseudotime inference. We found that smoothing process greatly increased the resolution of pseudotime tracks as compared to the ones directly inferred from the raw UMI counts (data not shown). Pseudotime inference was performed with default parameters according to the user manual (http://cole-trapnell-lab.github.io/monocle-release/docs/): 1) Set “negbinomial.size()” for expression distribution, and estimated size factors and dispersions. 2) Select genes detected among at least 5% of input cells to project cells to 2D space using “DDRTree” method. 3) Order cells and visualize pseudotime tracks as shown in Supplementary Fig. 1e and 2e. The ascending order of pseudotime values was consistent to the pattern of marker genes during spermatogenesis for both human and mouse (data not shown).

### Cell fate prediction with ‘RNA velocity’

We used the R package ‘velocyto.R’ (version 0.6) to estimate RNA velocity according to the standard procedures^26^. The RNA velocity estimation involves three separate counts matrices: intronic UMIs (nmat), exonic UMIs (emat), and the optional intron-exon spanning matrix (spmat). These matrices were generated by the ‘dropEst’ pipeline (version 0.7.1, https://github.com/hms-dbmi/dropEst). Briefly,1) The raw sequencing reads were tagged by droptag with the default ‘inDrop v1&v2’ configuration file except here that the ‘r1_rc_length’ was set as 3. 2) The tagged reads were mapped to the reference genomes (here human GRCh38 and mouse GRCm38) using STAR (version 2.5.3a) with default settings. 3) The alignments were processed by ‘dropEst’ with gene annotation GTF file (Ensembl release 90) and the default settings except here the ‘--merge-barcodes’ option was additionally called as suggested in the standard procedure. We followed the velocyto.R manual (https://github.com/velocyto-team/velocyto.R) which used emat and nmat to estimate and visualize RNA velocity. With predefined cell stage, we performed gene filtering with the parameter “min.max.cluster.average” set to 0.1 and 0.03 for emat and nmat, respectively. RNA velocity was estimated with the default settings except the parameters ‘kCells’ and ‘fit.quantile’ which were set as 3 and 0.05, respectively. RNA velocity field was visualized on a separate PCA embedding as shown in Fig. 1c for human germ cells, and in Supplementary Fig. 2a for mouse germ cells, respectively.

### Conservation and divergence analysis of human-mouse spermatogenesis

Following identifying the human and mouse spermatogenic cells separately, human-mouse spermatogenesis comparison was performed on genes which have one-to-one orthologues between human and mouse. Human-mouse one-to-one orthologous gene pair list was downloaded from Mouse Genome Informatics (MGI)-Vertebrate Homology (http://www.informatics.jax.org/homology.shtml). After filtering, 17,012 one-to-one orthologues genes were selected for integrating the human and mouse spermatogenic cells. Joint PCA was performed by selecting dynamically expressed genes using integrated gene expression matrix. In total, 1,124 genes were selected to perform joint PCA, as the results shown in Supplementary Fig. 2f-h. Top 20 genes contributing most to PC2 from both ends, which separated human and mouse species-specific signatures, were selected and plotted as shown in Supplementary Fig. 2i.

### Gene clustering

Gene clustering was performed on a collapsed expression matrix of genes-by-spermatogenic clusters. First, we defined the set of unexpressed genes by having expression (minimum of 1 UMI count per cell) in at least 5 single cells from the scRNA-seq data, or additionally, according to the specified parameter in Supplementary Fig. 3c by having a minimum expression level (mean UMI count for a stage as at least 0.1) in any give spermatogenic stage. The genes pass such criteria were defined as expressed genes. Expressed genes were then clustered by k-means algorithm, with k spread from 2 to 10, as shown in Supplementary Fig. 3c. Through interpreting the results, k=5 was chosen to display the gene clusters as it reflects the overall gene expression dynamics during spermatogenesis. Throughout the project we used gene clusters defined from germ cells from both biological and technical replicates, except for Supplementary Fig. 3a-b where we defined gene clusters from the two donors independently for sensitivity analysis.

The expressed genes were additionally clustered by their expression level, as used in the Fig. 4. The average expression level (UMI counts) across the spermatogenic cell clusters were used as input. To assign groups based on expression levels, we binned the genes by expression level into 9 groups:

Group 1: unexpressed;

Group 2: −inf < log_2_(UMI^mean^) ≤ −8;

Group 3: −8 < log_2_(UMI^mean^) ≤ −6;

Group 4: −6 < log_2_(UMI^mean^) ≤ −4;

Group 5: −4 < log_2_(UMI^mean^) ≤ −2;

Group 6: −2 < log_2_(UMI^mean^) ≤ 0;

Group 7: 0 < log_2_(UMI^mean^) ≤ 2;

Group 8: 2 < log_2_(UMI^mean^) ≤ 4;

Group 9: 4 < log_2_(UMI^mean^), highly expressed.

In addition, for modeling the germline variant levels versus expression level, the expression level was further binned into smaller groups. Specifically, log_2_(UMI^mean^) expression level between -8 and 4 were evenly binned into 100 expression level stages, and the genes within each expression level stage were isolated for calculating the germline variants levels and confidence intervals.

### Human and mouse germline variants

Human and mouse germline variations were downloaded from the Ensembl release 91 FTP site (ftp://ftp.ensembl.org/pub/release-91/variation/vcf/homo_sapiens/homo_sapiens.vcf.gz and ftp://ftp.ensembl.org/pub/release-91/variation/vcf/mus_musculus/mus_musculus.vcf.gz, respectively). We selected variants from dbSNP_150 and used BEDOPS together with custom Bash scripts to associate them with gene body, upstream 5kb, downstream 5kb genomic regions, and in addition, with the coding sequences and intron regions within the gene body. The gene body region was defined as the genomic interval between the gene start site and gene end site annotated in GTF file (Ensembl release 91). Because genes may have multiple different isoforms of transcripts with slightly different coding sequences, we broadly defined the genomic coding sequence regions as covered by coding sequences of any isoform mRNA. Introns was defined as regions where no coverage by coding sequences of any isoform mRNA. Moreover, we removed splicing consensus sequences – 6 bases on the 5’ end (splicing donor region) and 3 bases on the 3’ end (splicing acceptor region) – according to the gene orientation. With this strategy, we selected the intron regions with the least selection pressure. Upstream and downstream 5kb region was defined according to gene body region and with reference to gene orientation information. We classified the variants into the six mutation classes: (A>T/T>A; A>G/T>C; T>G/A>C; C>T/G>A; G>T/C>A; C>G/G>C). Each variant was then further distinguished in terms of the coding and the template strands, as previously introduced^45^. Then asymmetry score between the germline variants on the coding strand and template strand of each gene was calculated by log_2_(Var_coding_/Var_template_). The same procedures were also performed on upstream and downstream genomic regions, with the strand specificity (coding strand versus template strand) being assigned in consistent with the associated genes.

The germline mutation rates of the coding and the template stands were calculated by normalizing to a length of 1kb. Specifically, for germline mutations in total, the mutation rates were calculated as the sum of all germline short variants normalized to a length of 1kb. For specific base substitution mutation type, the mutation rates were calculated as the number of specific mutation type normalized to 1kb of the reference base type.

### Analyzing germline variants by gene family

Human gene family annotations were downloaded from the HUGO Gene Nomenclature Committee (https://www.genenames.org/data/genegroup/#!/). In total, 27 families contain more than 100 gene members. These families include: ‘Ankyrin repeat domain containing (ANKRD)’, ‘Armadillo-like helical domain containing (ARMH)’, ‘Basic helix-loop-helix proteins (BHLH)’, ‘BTB domain containing (BTBD)’, ‘Cadherins’, ‘CD molecules (CD)’, ‘EF-hand domain containing’, ‘Fibronectin type III domain containing’, ‘GPCR, Class A rhodopsin-like(excluding ORs)’, ‘GPCR, Class A rhodopsin-like(Olfactory receptors)’, ‘Heat shock proteins’, ‘Helicases’, ‘Histones’, ‘Homeoboxes’, ‘Immunoglobulin superfamily domain containing’, channels’, ‘PDZ domain containing (PDZ)’, ‘PHD finger proteins’, ‘Pleckstrin homology domain containing (PLEKH)’, ‘Ras small GTPases superfamily’, ‘Ring finger proteins’, ‘RNA binding motif containing (RBM)’, ‘Solute carriers (SLC)’, ‘WD repeat domain containing (WDR)’, ‘Zinc fingers C2H2-type’, ‘Zinc fingers - other’, ‘T cell receptor gene’. We further selected the gene families by having at least 10 gene members in both expressed and unexpressed categories, as defined above. This led to the selection of 10 gene families as shown in Fig. 2c. Levels of germline variant were calculated for each gene category of each family.

### Analyzing germline variants by GTEx expression profiles

The Genotype-Tissue Expression (GTEx) gene expression profiles used in Fig. 1e (release V7) across all the 53 tissue/organ/cell samples were downloaded from the GTEx Portal with release V7 (https://gtexportal.org/home/datasets/). We used the expression profiles containing the median TPM by tissue (GTEx_Analysis_2016-01-15_v7_RNASeQCv1.1.8_gene_median_tpm.gct.gz). To distinguish the expressed genes out of the unexpressed protein-coding genes for each tissue, we set the cutoff as 0.1 median TPM value as given from the GTEx Portal. For each tissue, a gene was defined as expressed if the expression level was ≥ 0.1, otherwise it was defined as unexpressed. Average germline variants associating with each gene category for each tissue was then calculated and the ratio was further calculated between the unexpressed gene category versus the expressed category. These ratios were plotted as shown in Fig. 2e. Z-scores were calculated on these ratios and indicated in the plot.

### Gene divergence datasets

The sequence divergence datasets of human to apes were downloaded from Ensembl release 91. Percent divergences in Fig. 5b and Supplementary Fig. 9f were calculated as: Divergence = 100% - Identity (human to other apes). dN and dS values were also retrieved from Ensembl and we excluded genes zero dN or dS. The mean values shown in Fig.5 and S9 were computed on non-outlier values, where an outlier value is defined as more than three scaled median absolute deviations (MAD) away from the median. For a set of divergence or dN/dS values made up N genes, MAD is defined as: MAD = median (|A*i* -median(A)|), for *i* = 1,2,…,N.

## Statistical Analysis

Statistical significance was computed by the Mann-Whitney U test (or rank-sum test) to test whether two groups of genes have distinct value distributions. Error bars of represents 99% percent confidence intervals, calculated by bootstrap methods sampling for 10,000 times.

## Data and code

The single cell RNA-seq sequencing results were deposited to NCBI GEO database with the accession code GSE125372.

---

**Supplementary Fig. 1.**
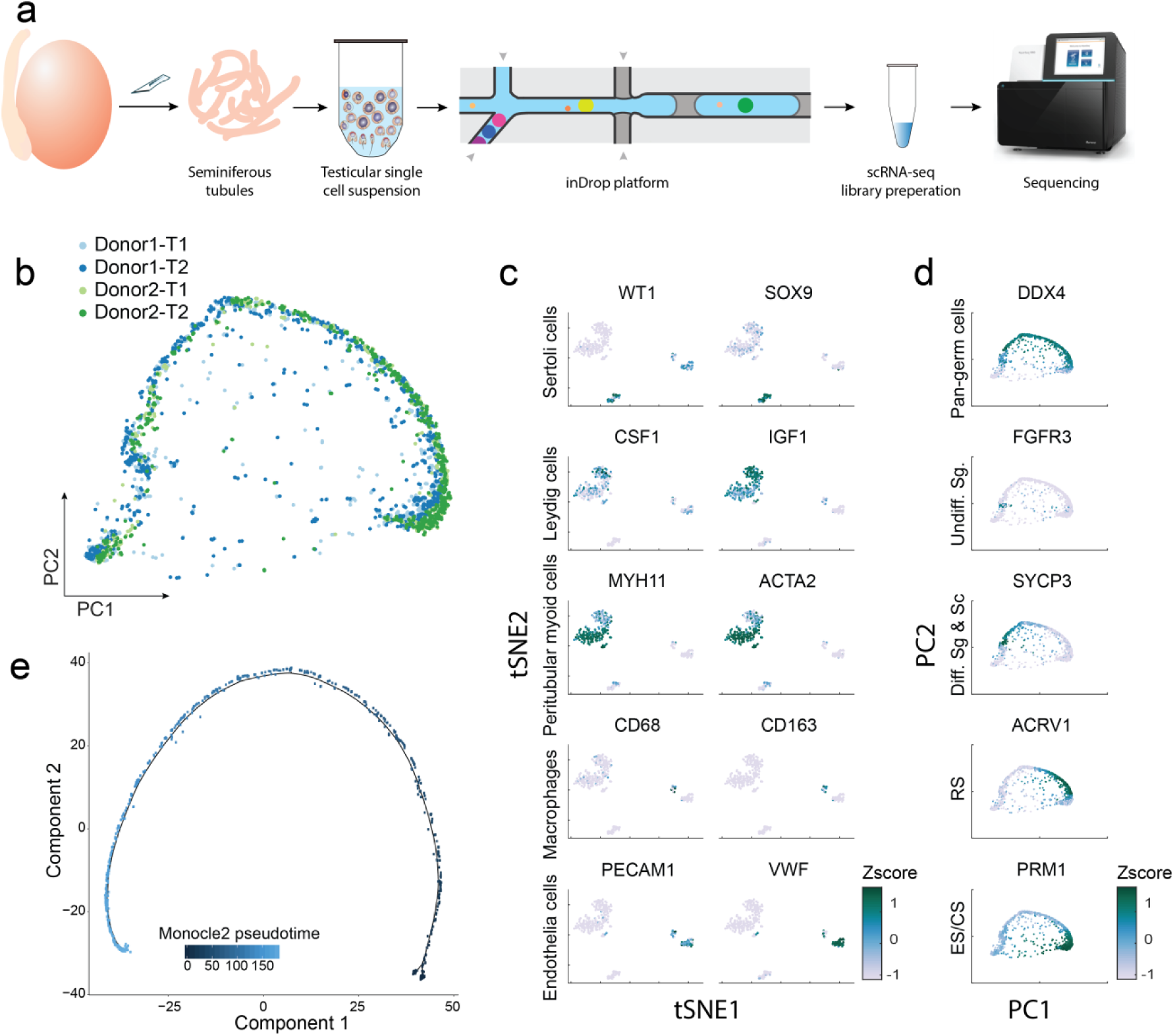
Single-cell transcriptomic analysis of human spermatogenesis. **a**, Schematic of single-cell RNA-seq of human testis samples with the inDrop microfluidics platform (Methods). **b**, Same PCA as in Figure 1B for the testicular cells across different specimen donors and technical replicates. **c-d**, Determining the identities of testicular somatic cells (**c**) and the developmental program of spermatogenic cells (**d**) (Methods). **e**, Human spermatogenic cell pseudotime defined by Monocle2 algorithm.

**Supplementary Fig. 2.**
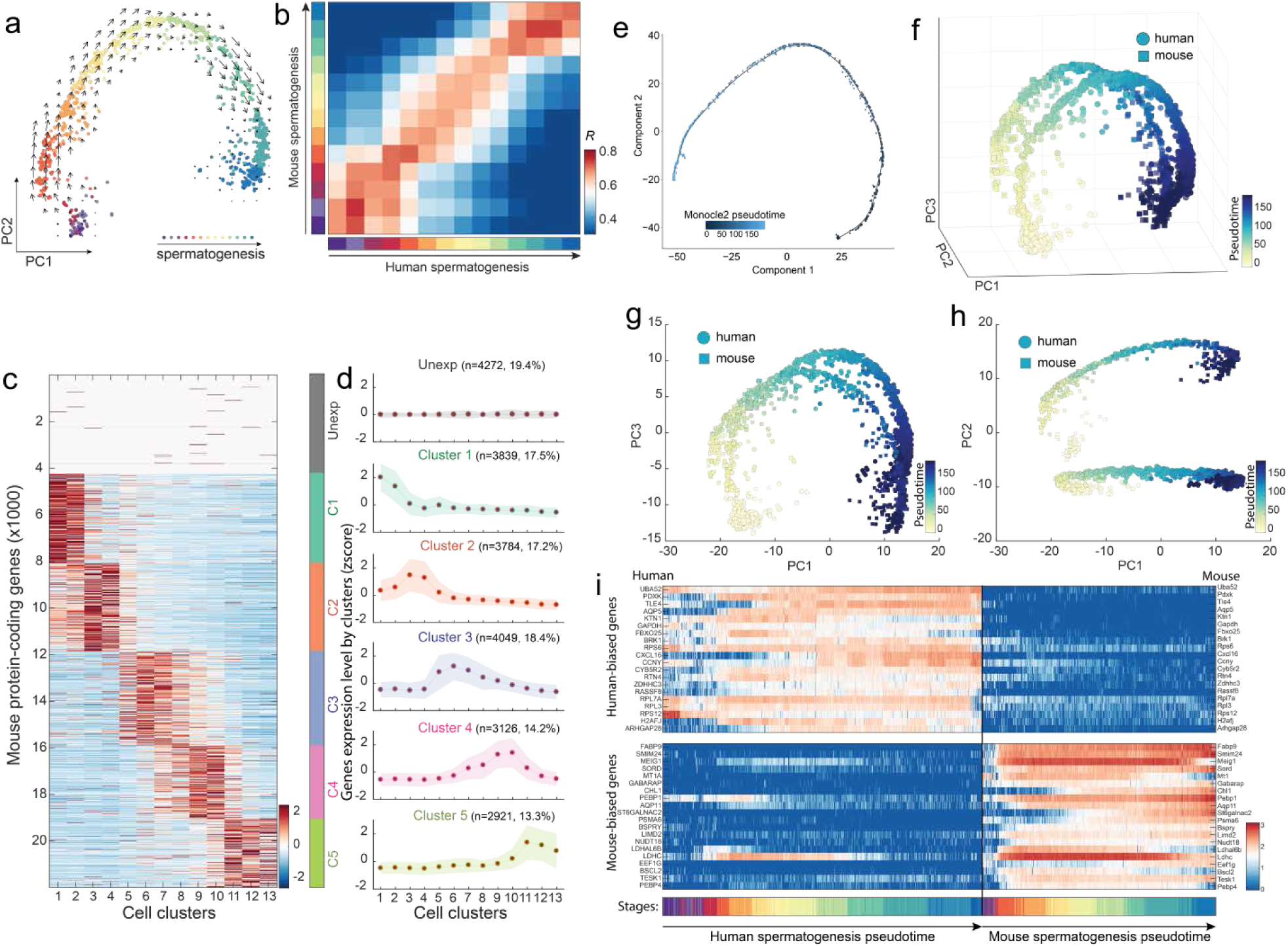
Single-cell transcriptomic analysis of mouse spermatogenesis comparison with human spermatogenesis. **a**, Principal components analysis on the spermatogenic-complement of the single-cell data. Arrows indicate the RNA velocity algorithm predicted developmental trajectory of mouse spermatogenesis. **b**, Correlation coefficients between human and mouse spermatogenic stages. **c**, Heatmap of all mouse proteins-coding genes clustered expression patterns. **d**, Expression profiles of mouse gene sets clustered by k-means clustering. **e**, Monocle2-ordering of human spermatogenic cells. **f**-**h**, Principal components analysis of all human and mouse spermatogenic cells mixed together. **i**, Expression heatmap of genes with highly divergent expression pattern between human and mouse spermatogenesis.

**Supplementary Fig. 3.**
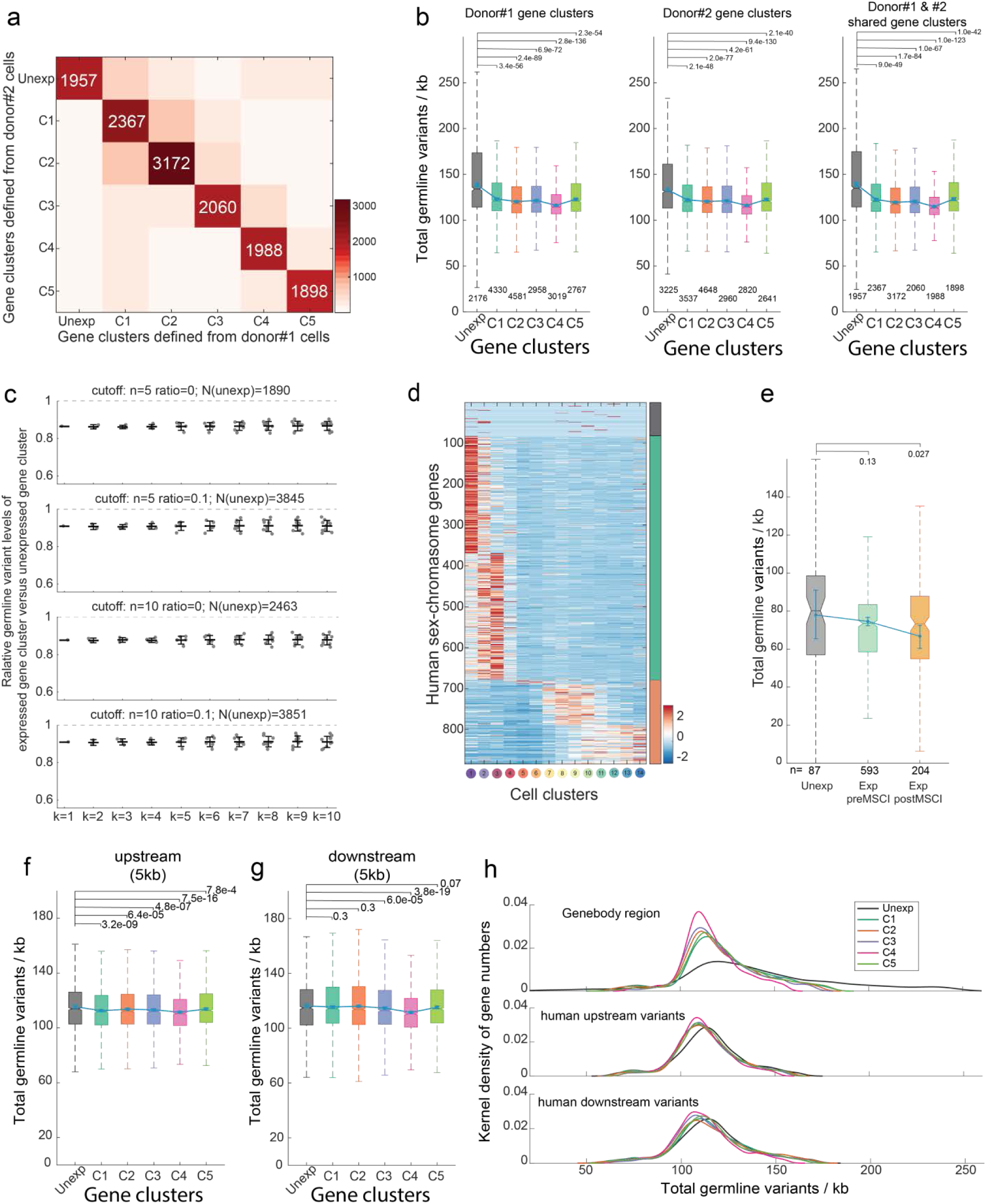
Control and sensitivity tests for the reduced germline mutation rates in the germ cell-expressed genes. **a**, Comparison of gene clusters determined by donor 1 germ cells and by donor 2 germ cells, respectively (Methods). The colors/numbers indicate the size of the cluster intersections. **b**, Same as in Figure 2b but using the gene clusters determined only by donor #1 germ cells (left), by donor #2 germ cells (middle) or by the intersection of genes of each cluster across donors (right). **c**, Ratios of germline mutation levels in the expressed genes versus the unexpressed genes. Each dot represents a specific ratio of germline mutation levels according to the corresponding expressed genes versus the unexpressed genes. The plot shows the ratios for k-means clustering of expressed genes with a range of k values. The four plots, show the results for a range of cutoffs used to determine the unexpressed gene cluster, generating different numbers (N) of unexpressed genes for sensitivity analysis. ‘cutoff n’ indicates the minimum number of cells expressing a given gene, and ‘ratio’ indicates the minimum expression level (average UMI) of a given gene in any one of the spermatogenesis cell clusters. **d**, Heatmap of human sex chromosome genes grouped into unexpressed genes, pre-meiotic sex chromosome inactivation (pre-MSCI) genes and post-MSCI genes. **e**, Human germline mutation rates across the sex chromosome gene clusters defined in (**d**). **f-g**, Germline mutation rates in both upstream 5kb (**f**) and downstream 5kb (**g**) of genes across clusters. **h**, Distributions of the germline mutation rates for each gene cluster defined in Fig. 1d, shown for the gene body region (top), upstream 5kb region (middle) and downstream 5kb region (bottom). Significance between mutation rates of expressed genes versus unexpressed genes is computed by the Mann-Whitney test with Bonferroni correction. Error bars indicate 99% confidence intervals calculated by bootstrap method with n=10,000.

**Supplementary Fig. 4.**
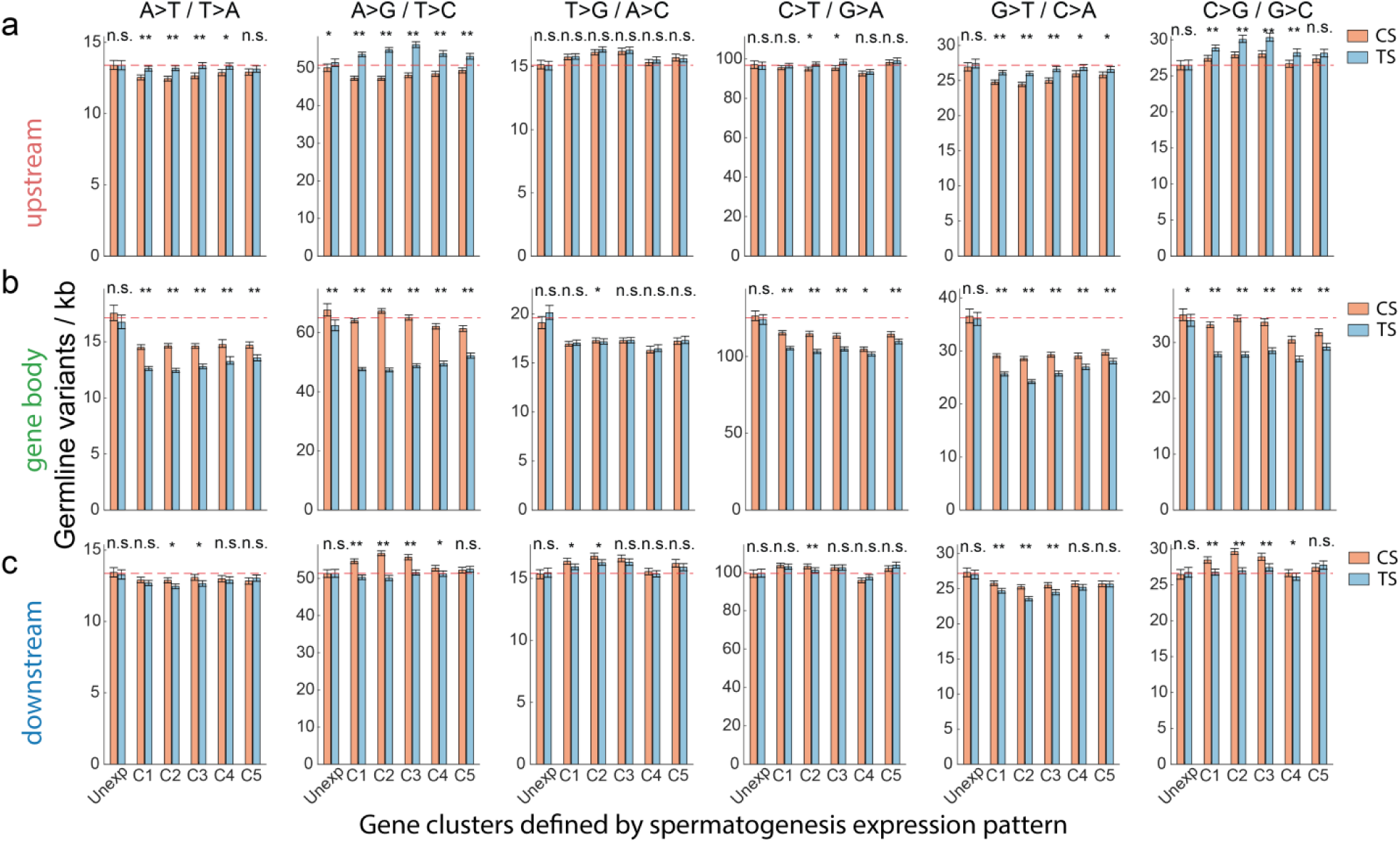
Germline mutation rates of gene body and flanking regions of all base-substitution mutation types. **a-c**, Germline mutation rates in the gene body region (**a**), upstream 5kb (**b**) and downstream 5kb (**c**). Dashed lines indicate the average level of mutations in unexpressed genes. Significance between mutation rates of coding strand versus that of template strand is computed by the Mann-Whitney test with Bonferroni correction. *, *P*<0.01; **, *P*<0.000001; n.s., not significant. Error bars indicate 99% confidence intervals calculated by bootstrap method with n=10,000.

**Supplementary Fig. 5.**
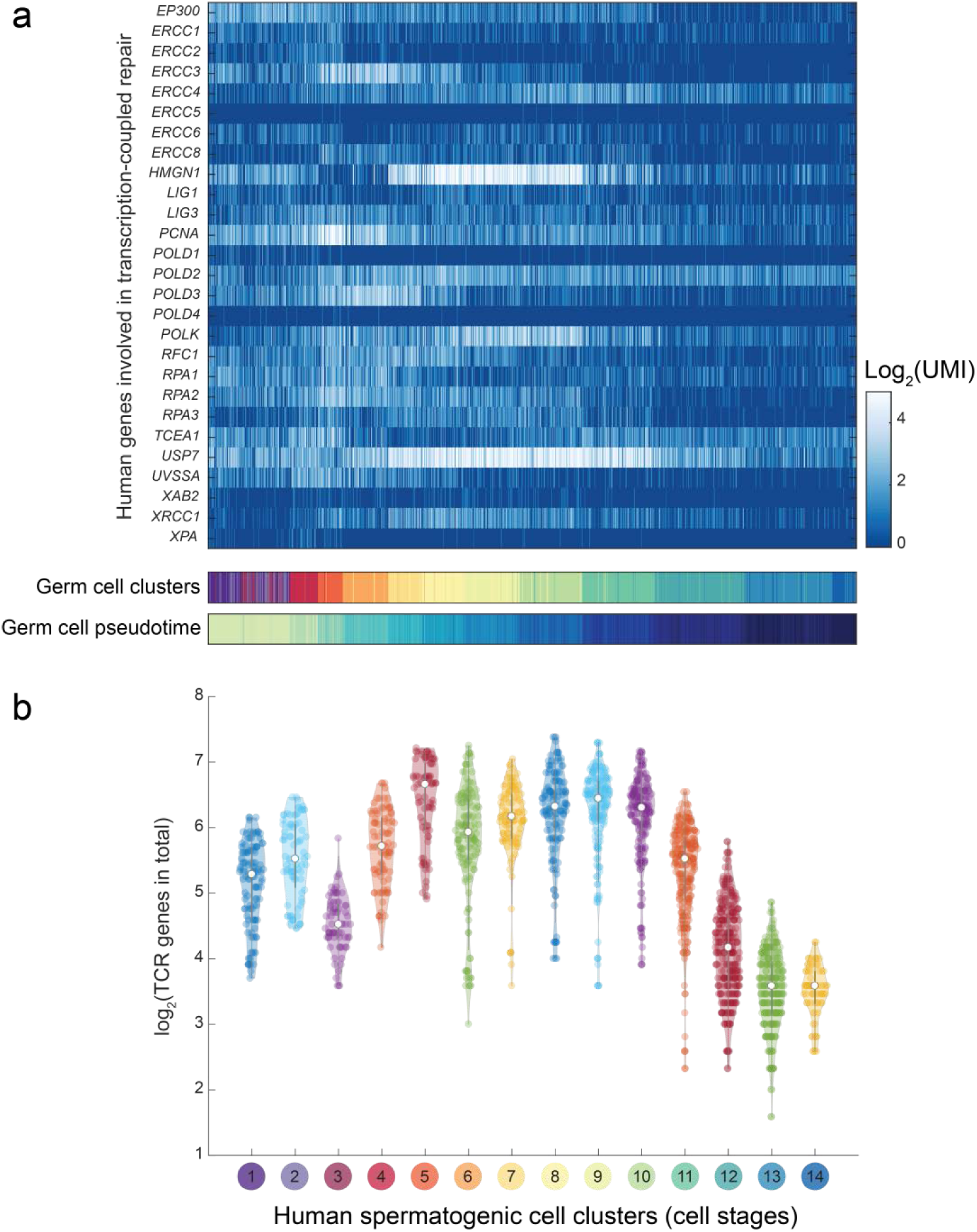
Gene expression profiles of genes involved in transcription-coupled repair (TCR). Gene expression levels of each TCR gene (**a**) and their sum (**b**) across all spermatogenic single cells are displayed, respectively.

**Supplementary Fig. 6.**
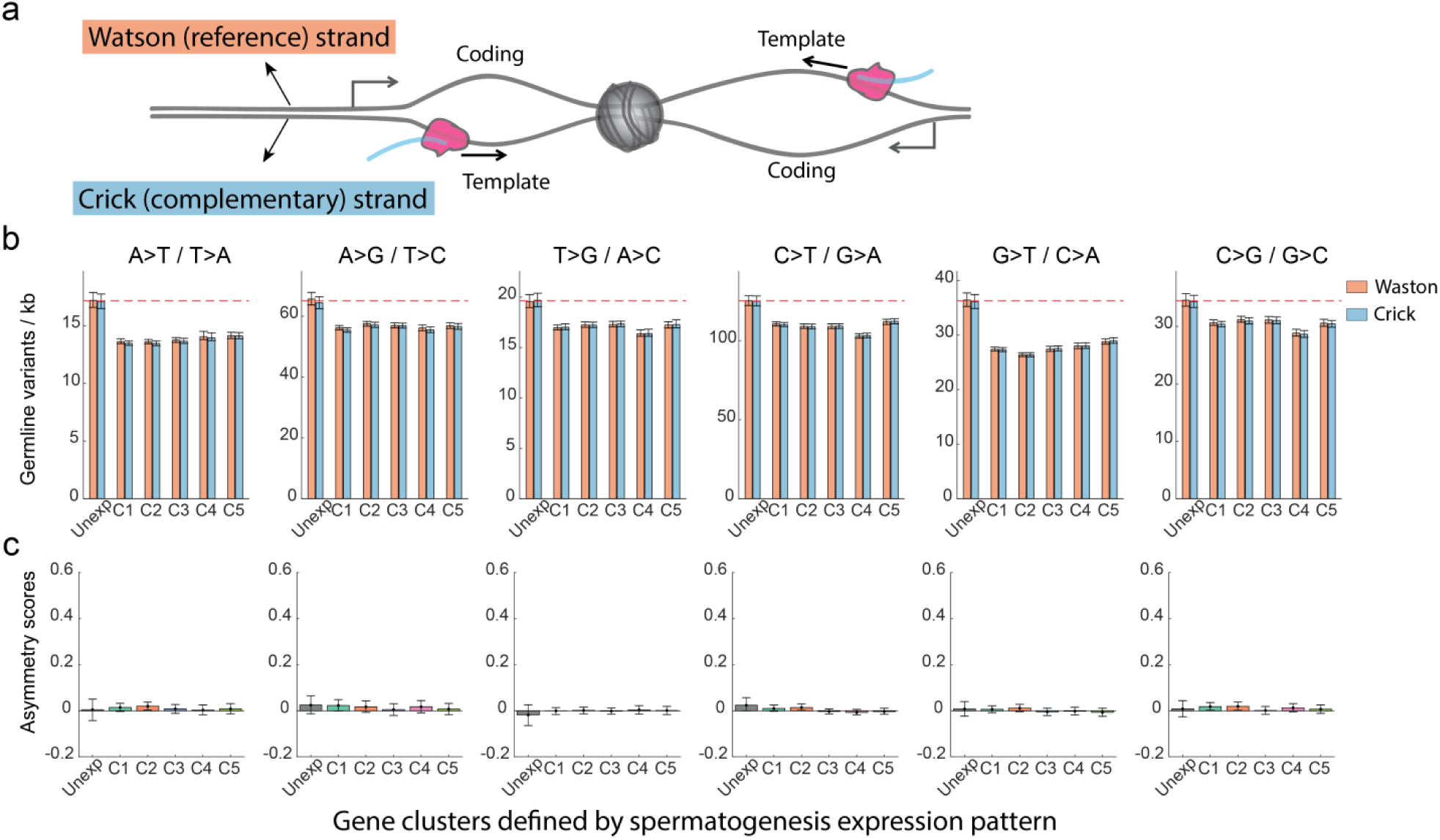
Mutation rate asymmetry is not detected between the Watson and Crick strands in expressed genes. **a**, Schematic of two neighboring genes, each on a different strand. Across the genome, genes are randomly disposed with respect to strand. **b-c**, Germline mutation rates (**b**) and asymmetry scores (**c**) of all base substitution mutation types across spermatogenesis expressed and unexpressed genes. Mutation rates and asymmetry scores were computed by distinguishing between the Watson and Crick strands, instead of coding and template strands (as shown in Fig. 3c and Supplementary Fig. 4). Dashed lines indicate the average level of mutations in unexpressed genes.

**Supplementary Fig. 7.**
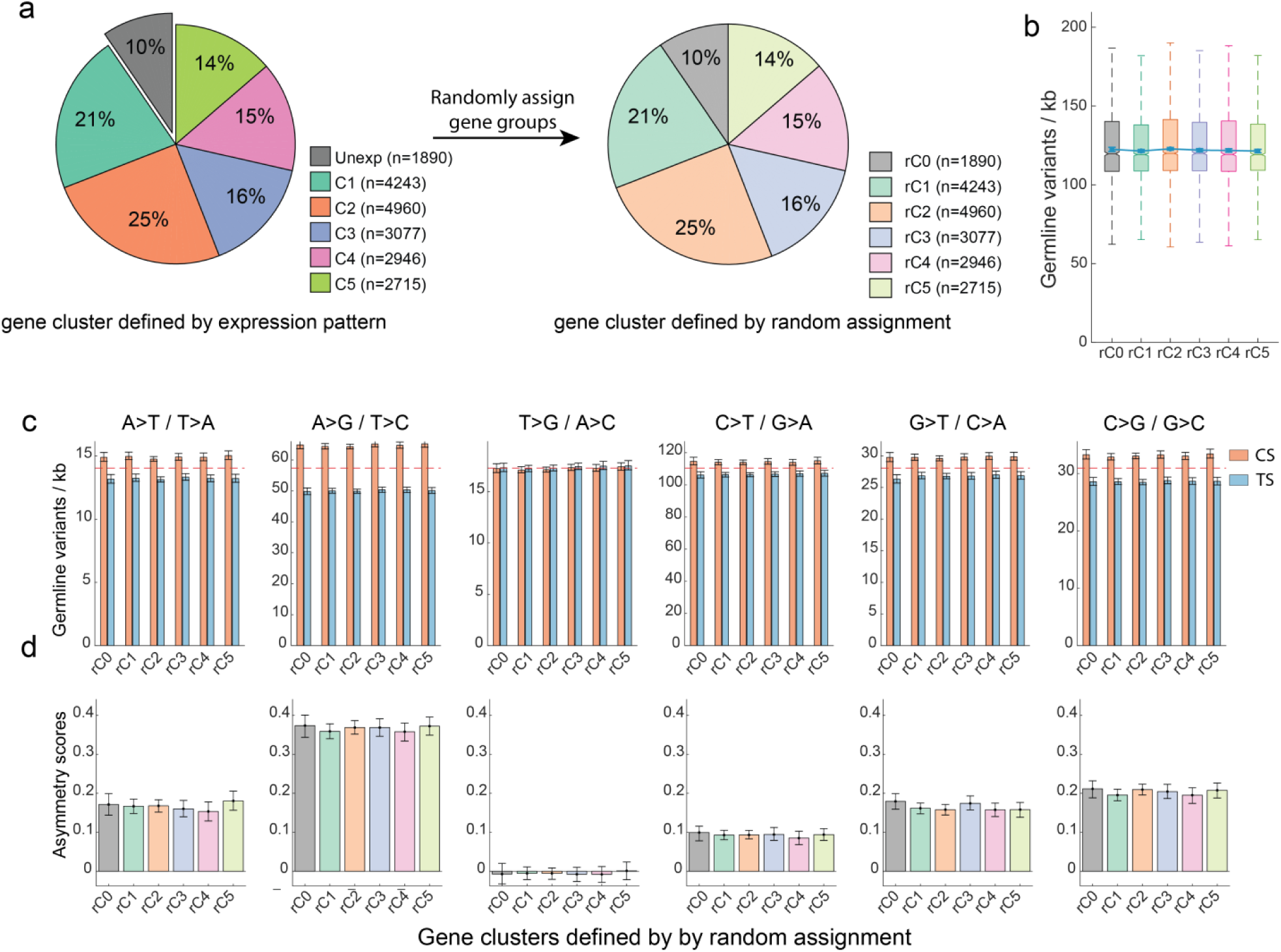
Shuffling gene assignments loses the mutation-level difference between expressed- and unexpressed genes. **a**, Shuffling gene group assignments. Genes assigned to all stages were shuffled, while maintaining the size of each group. **b**, Shuffling gene clustering loses the mutation-level differences between gene clusters. **c-d**, Germline mutation rates (**c**) and asymmetry scores (**d**) of all base substitution mutation types according to shuffled gene-grouping in (**a**). Mutation rates and asymmetry scores were computed by distinguishing between the coding and template strands (same as in Fig. 3c and Supplementary Fig. 4). Dashed lines indicate the average level of mutations in unexpressed genes.

**Supplementary Fig. 8.**
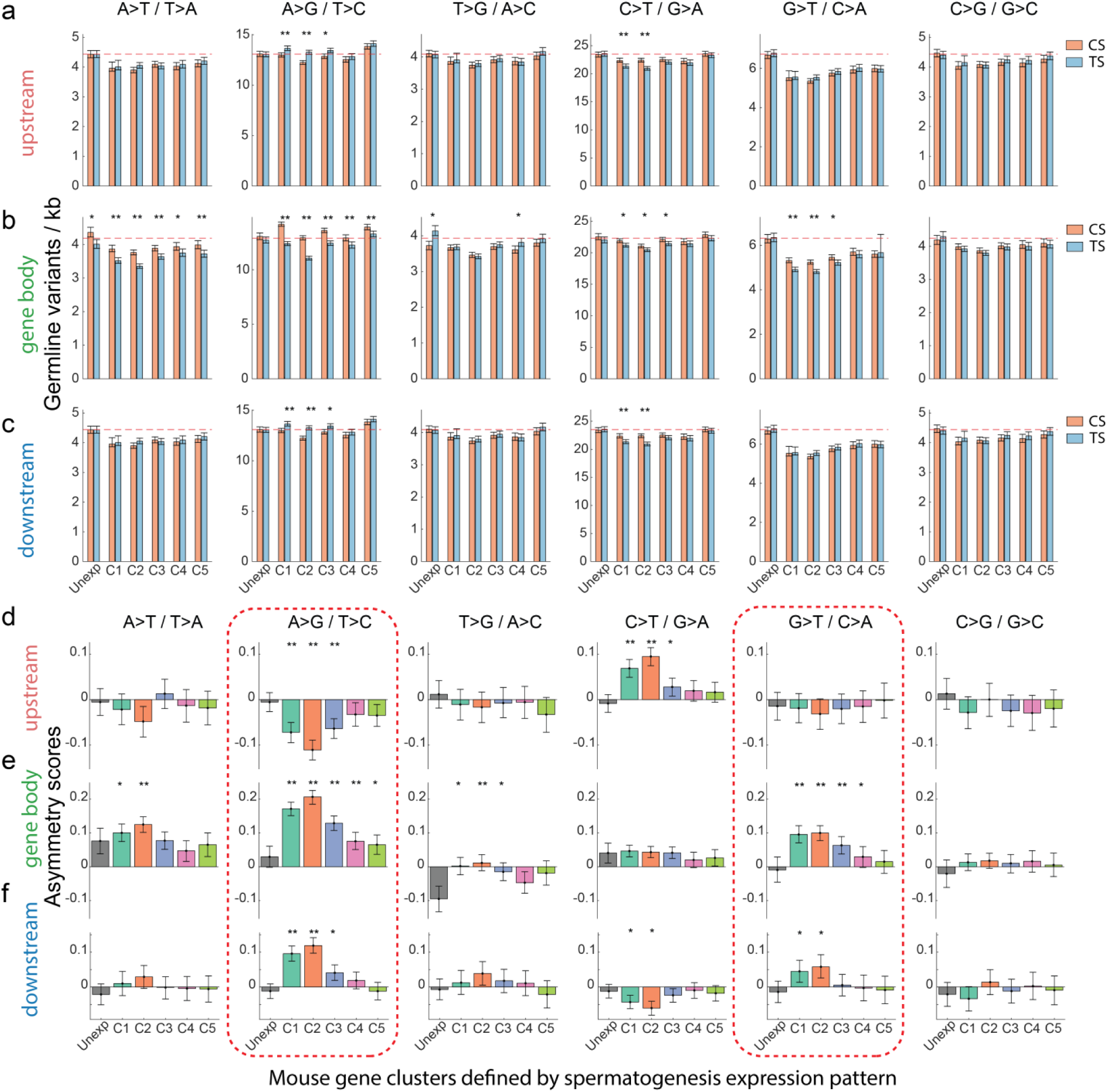
Mouse germline mutation rates and asymmetry scores of gene body and flanking regions of all base-substitution mutation types. **a-c**, Mouse germline mutation rates in the gene body region (**a**), upstream 5kb (**b**) and downstream 5kb (**c**). Dashed lines indicate the average level of mutations in unexpressed genes. **d-f**, Germline mutation asymmetry scores between coding and template strands in the upstream 5kb (**d**), gene body region (**e**) and downstream 5kb (**f**). Significance between germline variants on coding strand and template strand (**a**-**c**) or between the unexpressed category and the expressed gene categories (**d**-**f**) or is computed by the Mann-Whitney test with Bonferroni correction. *, *P*<0.01; **, *P*<0.000001; n.s., not significant. Error bars indicate 99% confidence intervals calculated by bootstrap method with n=10,000.

**Supplementary Fig. 9.**
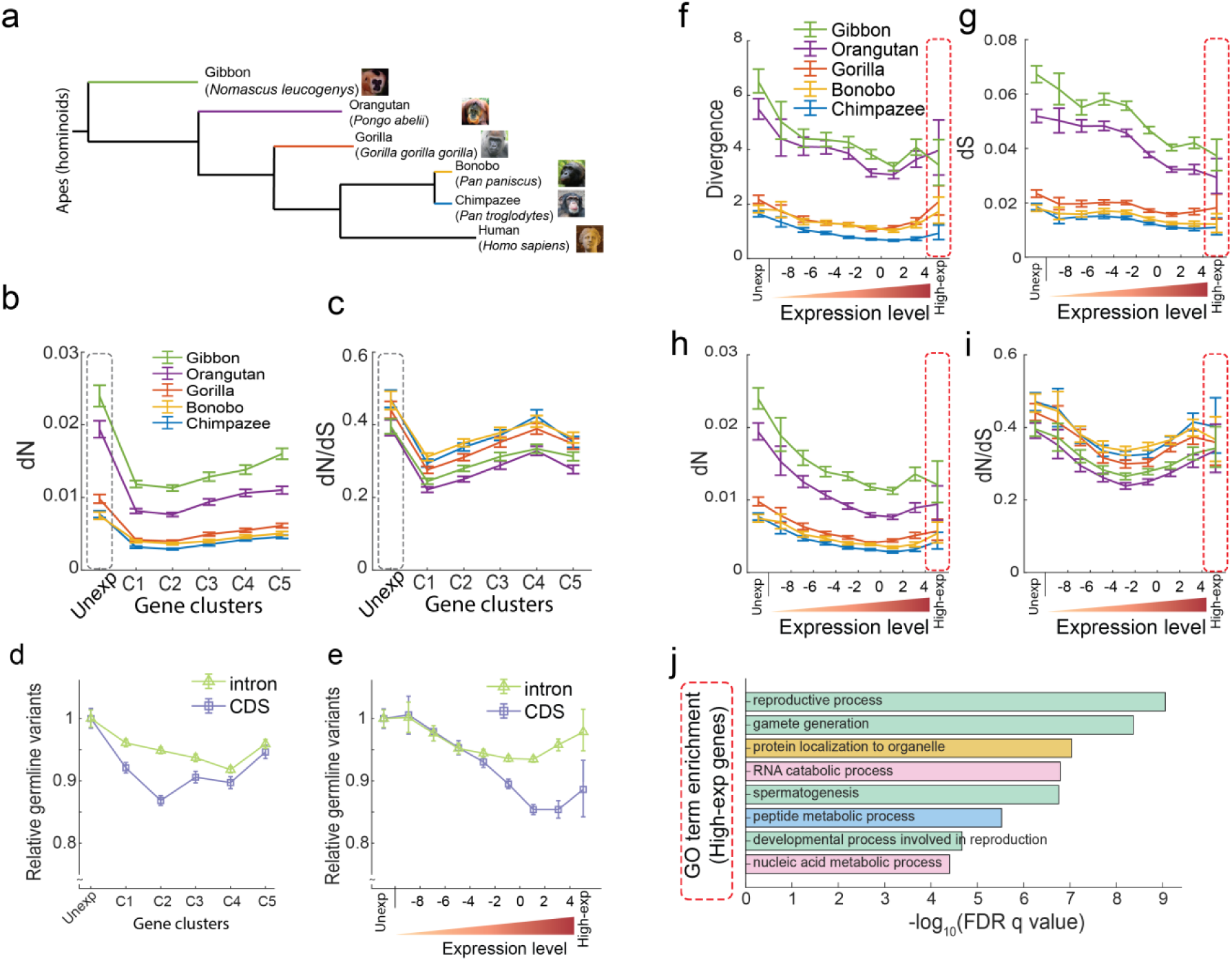
Evolutionary consequences of ‘transcriptional scanning’ across apes. **a**, Phylogenic tree of apes with sequenced genome data in Ensembl. **b-c**, dN (**b**) and dN/dS (**c**) values of human genes with their orthologues across apes, according to gene clusters defined from spermatogenesis expression. Grey dashed box highlights the unexpressed gene cluster. **d-e**, Relative germline mutations rates of intron regions and coding sequences according to gene expression-pattern clusters (**d**) and gene expression-level clusters (**e**). **f-i**, DNA divergence levels (**f**), dS scores (**g**), dN (**h**) and dN/dS (**i**) scores of human genes with their orthologues in the indicated apes, according to gene expression level categories. Red dashed box highlights the very highly expressed gene cluster. **j**, Gene ontology categories enriched in the set of genes that are very highly expressed during spermatogenesis.

**Supplementary Table 1.** Gene Ontology (GO) terms showing enrichment in the set of genes unexpressed in spermatogenesis. The GO term analysis was done by GOrilla^87^. ‘FDR q-value’ is the correction of p-values for multiple testing using the Benjamini and Hochberg method^58^. Enrichment (N, B, n, b) is defined as ‘Enrichment = (b/n) / (B/N)’. N, total number of genes; B, total number of genes associated with a specific GO term; n, number of genes in the input list; b, number of genes in the intersection. The highlighted GO terms are displayed in Fig. 5c.

| GO Term | Description | P-value | FDR q-value | Enrichment | N | B | n | b |
| --- | --- | --- | --- | --- | --- | --- | --- | --- |
| GO:0050907 | detection of chemical stimulus involved in sensory perception | 3.28E-171 | 5.07E-167 | 8.02 | 17883 | 406 | 1335 | 243 |
| GO:0009593 | detection of chemical stimulus | 1.07E-163 | 8.29E-160 | 7.51 | 17883 | 439 | 1335 | 246 |
| GO:0050911 | detection of chemical stimulus involved in sensory perception of smell | 2.31E-159 | 1.19E-155 | 8.27 | 17883 | 358 | 1335 | 221 |
| GO:0050906 | detection of stimulus involved in sensory perception | 3.18E-159 | 1.23E-155 | 7.24 | 17883 | 457 | 1335 | 247 |
| GO:0007186 | **G protein-coupled receptor signaling pathway** | 1.58E-144 | 4.87E-141 | 4.23 | 17883 | 1169 | 1335 | 369 |
| GO:0051606 | **detection of stimulus** | 4.21E-139 | 1.08E-135 | 5.84 | 17883 | 601 | 1335 | 262 |
| GO:0031424 | **keratinization** | 2.83E-51 | 6.25E-48 | 6.59 | 17883 | 179 | 1335 | 88 |
| GO:0007608 | sensory perception of smell | 3.95E-48 | 7.64E-45 | 7.65 | 17883 | 126 | 1335 | 72 |
| GO:0007606 | sensory perception of chemical stimulus | 1.90E-45 | 3.27E-42 | 6.57 | 17883 | 159 | 1335 | 78 |
| GO:0050896 | response to stimulus | 7.85E-41 | 1.21E-37 | 1.58 | 17883 | 5109 | 1335 | 602 |
| GO:0007165 | signal transduction | 4.67E-39 | 6.56E-36 | 1.62 | 17883 | 4480 | 1335 | 543 |
| GO:0006955 | **immune response** | 1.16E-25 | 1.50E-22 | 2.37 | 17883 | 905 | 1335 | 160 |
| GO:0007600 | **sensory perception** | 8.52E-25 | 1.01E-21 | 2.85 | 17883 | 526 | 1335 | 112 |
| GO:0006952 | **defense response** | 7.32E-22 | 8.08E-19 | 2.14 | 17883 | 1045 | 1335 | 167 |
| GO:0032501 | multicellular organismal process | 1.70E-21 | 1.75E-18 | 1.55 | 17883 | 3315 | 1335 | 384 |
| GO:0098542 | defense response to other organism | 6.13E-19 | 5.92E-16 | 2.8 | 17883 | 416 | 1335 | 87 |
| GO:0050877 | nervous system process | 4.34E-18 | 3.94E-15 | 2.12 | 17883 | 892 | 1335 | 141 |
| GO:0033141 | positive regulation of peptidyl-serine phosphorylation of STAT protein | 4.99E-18 | 4.28E-15 | 11.48 | 17883 | 21 | 1335 | 18 |
| GO:0033139 | regulation of peptidyl-serine phosphorylation of STAT protein | 1.09E-16 | 8.88E-14 | 10.48 | 17883 | 23 | 1335 | 18 |
| GO:0003008 | system process | 2.87E-16 | 2.22E-13 | 1.81 | 17883 | 1366 | 1335 | 185 |
| GO:0002323 | natural killer cell activation involved in immune response | 4.06E-16 | 2.99E-13 | 10.05 | 17883 | 24 | 1335 | 18 |
| GO:0042742 | defense response to bacterium | 5.65E-16 | 3.97E-13 | 3.32 | 17883 | 226 | 1335 | 56 |
| GO:0006959 | **humoral immune response** | 5.33E-15 | 3.59E-12 | 3.29 | 17883 | 216 | 1335 | 53 |
| GO:0051707 | response to other organism | 2.27E-14 | 1.46E-11 | 2.21 | 17883 | 613 | 1335 | 101 |
| GO:0001580 | detection of chemical stimulus involved in sensory perception of bitter taste | 3.14E-14 | 1.94E-11 | 7.21 | 17883 | 39 | 1335 | 21 |
| GO:0050912 | detection of chemical stimulus involved in sensory perception of taste | 1.06E-13 | 6.28E-11 | 6.55 | 17883 | 45 | 1335 | 22 |
| GO:0045087 | **innate immune response** | 6.95E-13 | 3.98E-10 | 2.38 | 17883 | 434 | 1335 | 77 |
| GO:0009617 | response to bacterium | 8.29E-13 | 4.58E-10 | 2.66 | 17883 | 312 | 1335 | 62 |
| GO:0043207 | response to external biotic stimulus | 1.55E-12 | 8.25E-10 | 1.93 | 17883 | 825 | 1335 | 119 |
| GO:0030101 | natural killer cell activation | 6.92E-12 | 3.56E-09 | 5.31 | 17883 | 58 | 1335 | 23 |
| GO:0002250 | adaptive immune response | 7.82E-12 | 3.90E-09 | 2.98 | 17883 | 211 | 1335 | 47 |
| GO:0009607 | response to biotic stimulus | 1.07E-11 | 5.16E-09 | 1.88 | 17883 | 849 | 1335 | 119 |
| GO:0007210 | serotonin receptor signaling pathway | 2.38E-10 | 1.11E-07 | 6.15 | 17883 | 37 | 1335 | 17 |
| GO:0002376 | immune system process | 6.19E-10 | 2.81E-07 | 1.48 | 17883 | 2014 | 1335 | 222 |
| GO:0042100 | B cell proliferation | 1.07E-09 | 4.72E-07 | 5.69 | 17883 | 40 | 1335 | 17 |
| GO:0010469 | regulation of signaling receptor activity | 1.78E-09 | 7.63E-07 | 1.99 | 17883 | 546 | 1335 | 81 |
| GO:0007187 | G protein-coupled receptor signaling pathway, coupled to cyclic nucleotide second messenger | 1.06E-08 | 4.42E-06 | 2.58 | 17883 | 218 | 1335 | 42 |
| GO:0043330 | response to exogenous dsRNA | 1.37E-08 | 5.57E-06 | 4.95 | 17883 | 46 | 1335 | 17 |
| GO:0050830 | defense response to Gram-positive bacterium | 1.45E-08 | 5.75E-06 | 3.56 | 17883 | 94 | 1335 | 25 |
| GO:0043331 | response to dsRNA | 1.71E-08 | 6.60E-06 | 4.64 | 17883 | 52 | 1335 | 18 |
| GO:0002286 | T cell activation involved in immune response | 1.71E-08 | 6.44E-06 | 4.64 | 17883 | 52 | 1335 | 18 |
| GO:0098664 | G protein-coupled serotonin receptor signaling pathway | 2.05E-08 | 7.53E-06 | 5.86 | 17883 | 32 | 1335 | 14 |
| GO:0009605 | response to external stimulus | 2.93E-08 | 1.05E-05 | 1.52 | 17883 | 1440 | 1335 | 163 |
| GO:0018149 | peptide cross-linking | 3.62E-08 | 1.27E-05 | 4.24 | 17883 | 60 | 1335 | 19 |
| GO:0050829 | defense response to Gram-negative bacterium | 5.48E-08 | 1.88E-05 | 3.68 | 17883 | 80 | 1335 | 22 |
| GO:0007218 | neuropeptide signaling pathway | 8.45E-08 | 2.84E-05 | 3.28 | 17883 | 102 | 1335 | 25 |
| GO:1904892 | regulation of STAT cascade | 1.37E-07 | 4.50E-05 | 2.98 | 17883 | 126 | 1335 | 28 |
| GO:0046425 | regulation of JAK-STAT cascade | 2.49E-07 | 8.01E-05 | 2.96 | 17883 | 122 | 1335 | 27 |

**Supplementary Table 2.** Gene Ontology terms showing enrichment in the set of genes that are highly-expressed throughout spermatogenesis. The GO term analysis was done in the same way as described in Supplementary Table 1.

| GO Term | Description | P-value | FDR<br>q-value | Enrichment | N | B | n | b |
| --- | --- | --- | --- | --- | --- | --- | --- | --- |
| GO:0006614 | SRP-dependent cotranslational protein targeting to membrane | 2.52E-14 | 3.89E-10 | 18.47 | 17883 | 88 | 154 | 14 |
| GO:0006613 | cotranslational protein targeting to membrane | 5.60E-14 | 4.33E-10 | 17.48 | 17883 | 93 | 154 | 14 |
| GO:0006413 | translational initiation | 6.33E-14 | 3.26E-10 | 13.37 | 17883 | 139 | 154 | 16 |
| GO:0000184 | nuclear-transcribed mRNA catabolic process, nonsense-mediated decay | 7.90E-14 | 3.05E-10 | 14.89 | 17883 | 117 | 154 | 15 |
| GO:0045047 | protein targeting to ER | 2.10E-13 | 6.49E-10 | 15.94 | 17883 | 102 | 154 | 14 |
| GO:0072599 | establishment of protein localization to endoplasmic reticulum | 3.63E-13 | 9.34E-10 | 15.34 | 17883 | 106 | 154 | 14 |
| GO:0022414 | **reproductive process** | 3.98E-13 | 8.79E-10 | 3.56 | 17883 | 1339 | 154 | 41 |
| GO:0072594 | establishment of protein localization to organelle | 1.49E-12 | 2.87E-09 | 6.76 | 17883 | 378 | 154 | 22 |
| GO:0048609 | multicellular organismal reproductive process | 1.49E-12 | 2.56E-09 | 5.47 | 17883 | 552 | 154 | 26 |
| GO:0070972 | protein localization to endoplasmic reticulum | 1.64E-12 | 2.53E-09 | 13.78 | 17883 | 118 | 154 | 14 |
| GO:0007276 | **gamete generation** | 3.05E-12 | 4.29E-09 | 6.15 | 17883 | 434 | 154 | 23 |
| GO:0000956 | nuclear-transcribed mRNA catabolic process | 6.50E-12 | 8.37E-09 | 9.94 | 17883 | 187 | 154 | 16 |
| GO:0006402 | mRNA catabolic process | 2.27E-11 | 2.70E-08 | 9.15 | 17883 | 203 | 154 | 16 |
| GO:0006612 | protein targeting to membrane | 4.86E-11 | 5.37E-08 | 10.77 | 17883 | 151 | 154 | 14 |
| GO:0033365 | **protein localization to organelle** | 8.96E-11 | 9.24E-08 | 4.74 | 17883 | 612 | 154 | 25 |
| GO:0006401 | **RNA catabolic process** | 1.68E-10 | 1.62E-07 | 8.01 | 17883 | 232 | 154 | 16 |
| GO:0007283 | **spermatogenesis** | 1.93E-10 | 1.75E-07 | 5.89 | 17883 | 394 | 154 | 20 |
| GO:0048232 | male gamete generation | 2.11E-10 | 1.81E-07 | 5.86 | 17883 | 396 | 154 | 20 |
| GO:0090150 | establishment of protein localization to membrane | 2.61E-10 | 2.12E-07 | 7.77 | 17883 | 239 | 154 | 16 |
| GO:0006605 | protein targeting | 1.44E-09 | 1.12E-06 | 6.39 | 17883 | 309 | 154 | 17 |
| GO:0006412 | translation | 1.56E-09 | 1.15E-06 | 8.29 | 17883 | 196 | 154 | 14 |
| GO:0022412 | cellular process involved in reproduction in multicellular organism | 1.84E-09 | 1.29E-06 | 6.29 | 17883 | 314 | 154 | 17 |
| GO:0006518 | **peptide metabolic process** | 4.46E-09 | 3.00E-06 | 5.93 | 17883 | 333 | 154 | 17 |
| GO:0043043 | peptide biosynthetic process | 5.82E-09 | 3.75E-06 | 7.49 | 17883 | 217 | 154 | 14 |
| GO:0034655 | nucleobase-containing compound catabolic process | 8.35E-09 | 5.16E-06 | 5.32 | 17883 | 393 | 154 | 18 |
| GO:0044772 | mitotic cell cycle phase transition | 1.29E-08 | 7.69E-06 | 7.04 | 17883 | 231 | 154 | 14 |
| GO:0044770 | cell cycle phase transition | 1.99E-08 | 1.14E-05 | 6.8 | 17883 | 239 | 154 | 14 |
| GO:0003006 | **developmental process involved in reproduction** | 3.88E-08 | 2.14E-05 | 4.12 | 17883 | 592 | 154 | 21 |
| GO:0046700 | heterocycle catabolic process | 4.56E-08 | 2.43E-05 | 4.76 | 17883 | 439 | 154 | 18 |
| GO:0044270 | cellular nitrogen compound catabolic process | 4.72E-08 | 2.43E-05 | 4.75 | 17883 | 440 | 154 | 18 |
| GO:0019439 | aromatic compound catabolic process | 7.32E-08 | 3.65E-05 | 4.61 | 17883 | 453 | 154 | 18 |
| GO:0090304 | **nucleic acid metabolic process** | 8.21E-08 | 3.97E-05 | 2.31 | 17883 | 2163 | 154 | 43 |
| GO:0034645 | cellular macromolecule<br>biosynthetic process | 9.17E-08 | 4.29E-05 | 3.16 | 17883 | 991 | 154 | 27 |
| GO:0000086 | G2/M transition of mitotic cell<br>cycle | 1.09E-07 | 4.96E-05 | 9.44 | 17883 | 123 | 154 | 10 |
| GO:0044839 | cell cycle G2/M phase transition | 1.27E-07 | 5.62E-05 | 9.29 | 17883 | 125 | 154 | 10 |
| GO:0043604 | amide biosynthetic process | 1.42E-07 | 6.11E-05 | 5.36 | 17883 | 325 | 154 | 15 |
| GO:0022402 | cell cycle process | 1.49E-07 | 6.22E-05 | 3.28 | 17883 | 885 | 154 | 25 |
| GO:1901361 | organic cyclic compound catabolic<br>process | 2.09E-07 | 8.49E-05 | 4.3 | 17883 | 486 | 154 | 18 |

## References

1. E. E. Schmidt, U. Schibler, High accumulation of components of the RNA polymerase II transcription machinery in rodent spermatids. Development. 121, 2373–2383 (1995).

2. D. S. Johnston et al., Stage-specific gene expression is a fundamental characteristic of rat spermatogenic cells and Sertoli cells. Proc. Natl. Acad. Sci. USA. 105, 8315–8320 (2008).

3. D. Brawand et al., The evolution of gene expression levels in mammalian organs. Nature. 478, 343–348 (2011).

4. M. Melé et al., The human transcriptome across tissues and individuals. Science (80-.). 348, 660–665 (2015).

5. M. Soumillon et al., Cellular source and mechanisms of high transcriptome complexity in the mammalian testis. Cell Rep. 3, 2179–2190 (2013).

6. D. Djureinovic et al., The human testis-specific proteome defined by transcriptomics and antibody-based profiling. Mol. Hum. Reprod. 20, 476–488 (2014).

7. E. E. Schmidt, Transcriptional promiscuity in testes. Curr. Biol. 6, 768–769 (1996).

8. K. C. Kleene, A possible meiotic function of the peculiar patterns of gene expression in mammalian spermatogenic cells. Mech. Dev. 106, 3–23 (2001).

9. B. B. Lake et al., Neuronal subtypes and diversity revealed by single-nucleus RNA sequencing of the human brain. Science (80-.). 352, 1586–1590 (2016).

10. H. Miyata et al., Genome engineering uncovers 54 evolutionarily conserved and testis-enriched genes that are not required for male fertility in mice. Proc. Natl. Acad. Sci. USA. 113, 7704–7710 (2016).

11. D. Wang et al., A deep proteome and transcriptome abundance atlas of 29 healthy human tissues. BioRxiv 3. (2018), doi:10.1101/357137.

12. K. C. Kleene, Patterns, mechanisms, and functions of translation regulation in mammalian spermatogenic cells. Cytogenet Genome Res. 103, 217–224 (2003).

13. C. Rathke, W. M. Baarends, S. Awe, R. Renkawitz-Pohl, Chromatin dynamics during spermiogenesis. Biochim. Biophys. Acta. 1839, 155–168 (2014).

14. A. Necsulea, H. Kaessmann, Evolutionary dynamics of coding and non-coding transcriptomes. Nat. Rev. Genet. 15, 734–748 (2014).

15. C. Naro et al., An Orchestrated Intron Retention Program in Meiosis Controls Timely Usage of Transcripts during Germ Cell Differentiation. Dev. Cell. 41, 82–93.e4 (2017).

16. M. Lynch, G. K. Marinov, The bioenergetic costs of a gene. Proc. Natl. Acad. Sci. USA. 112, 15690–15695 (2015).

17. L. Huang, Z. Yuan, J. Yu, T. Zhou, Fundamental principles of energy consumption for gene expression. Chaos. 25, 123101 (2015).

18. I. Frumkin et al., Gene Architectures that Minimize Cost of Gene Expression. Mol. Cell. 65, 142–153 (2017).

19. P. C. Hanawalt, G. Spivak, Transcription-coupled DNA repair: two decades of progress and surprises. Nat. Rev. Mol. Cell Biol. 9, 958–970 (2008).

20. W. Vermeulen, M. Fousteri, Mammalian transcription-coupled excision repair. Cold Spring Harb. Perspect. Biol. 5, a012625 (2013).

21. A. Werner, M. J. Piatek, J. S. Mattick, Transpositional shuffling and quality control in male germ cells to enhance evolution of complex organisms. Ann. N. Y. Acad. Sci. 1341, 156–163 (2015).

22. M. F. Flajnik, M. Kasahara, Origin and evolution of the adaptive immune system: genetic events and selective pressures. Nat. Rev. Genet. 11, 47–59 (2010).

23. T. Boehm, Evolution of vertebrate immunity. Curr. Biol. 22, R722–32 (2012).

24. R. S. Singh, J. Xu, R. J. Kulathinal, Rapidly evolving genes and genetic systems (books.google.com, 2012).

25. S. Jinks-Robertson, A. S. Bhagwat, Transcription-associated mutagenesis. Annu. Rev. Genet. 48, 341–359 (2014).

26. G. La Manno et al., RNA velocity of single cells. Nature. 560, 494–498 (2018).

27. A. M. Klein et al., Droplet barcoding for single-cell transcriptomics applied to embryonic stem cells. Cell. 161, 1187–1201 (2015).

28. M. Kanatsu-Shinohara, T. Shinohara, Spermatogonial stem cell self-renewal and development. Annu. Rev. Cell Dev. Biol. 29, 163–187 (2013).

29. S. S. Hammoud et al., Chromatin and transcription transitions of mammalian adult germline stem cells and spermatogenesis. Cell Stem Cell. 15, 239–253 (2014).

30. R. Sharma, A. Agarwal, in Sperm Chromatin, A. Zini, A. Agarwal, Eds. (Springer New York, New York, NY, 2011), pp. 19–44.

31. X. Qiu et al., Reversed graph embedding resolves complex single-cell trajectories. Nat. Methods. 14, 979–982 (2017).

32. D. Paoli et al., Sperm glyceraldehyde 3-phosphate dehydrogenase gene expression in asthenozoospermic spermatozoa. Asian J Androl. 19, 409–413 (2017).

33. V. Selvaraj et al., Mice lacking FABP9/PERF15 develop sperm head abnormalities but are fertile. Dev. Biol. 348, 177–189 (2010).

34. G. Frenette, M. Thabet, R. Sullivan, Polyol pathway in human epididymis and semen. J Androl. 27, 233–239 (2006).

35. GTEx Consortium, Human genomics. The Genotype-Tissue Expression (GTEx) pilot analysis: multitissue gene regulation in humans. Science (80-.). 348, 648–660 (2015).

36. D. R. Zerbino et al., Ensembl 2018. Nucleic Acids Res. 46, D754–D761 (2018).

37. K. D. Makova, W.-H. Li, Strong male-driven evolution of DNA sequences in humans and apes. Nature. 416, 624–626 (2002).

38. C. D. Campbell, E. E. Eichler, Properties and rates of germline mutations in humans. Trends Genet. 29, 575–584 (2013).

39. R. Acuna-Hidalgo, J. A. Veltman, A. Hoischen, New insights into the generation and role of de novo mutations in health and disease. Genome Biol. 17, 241 (2016).

40. A. Tubbs, A. Nussenzweig, Endogenous DNA damage as a source of genomic instability in cancer. Cell. 168, 644–656 (2017).

41. M. Nei, Y. Suzuki, M. Nozawa, The neutral theory of molecular evolution in the genomic era. Annu. Rev. Genomics. Hum. Genet. 11, 265–289 (2010).

42. G. Xu et al., Nucleotide excision repair activity varies among murine spermatogenic cell types. Biol. Reprod. 73, 123–130 (2005).

43. C. Chen, H. Qi, Y. Shen, J. Pickrell, M. Przeworski, Contrasting determinants of mutation rates in germline and soma. Genetics. 207, 255–267 (2017).

44. K. A. Gray, R. L. Seal, S. Tweedie, M. W. Wright, E. A. Bruford, A review of the new HGNC gene family resource. Hum. Genomics. 10, 6 (2016).

45. N. J. Haradhvala et al., Mutational strand asymmetries in cancer genomes reveal mechanisms of DNA damage and repair. Cell. 164, 538–549 (2016).

46. P. Green et al., Transcription-associated mutational asymmetry in mammalian evolution. Nat. Genet. 33, 514–517 (2003).

47. C. F. Mugal, H.-H. von Grünberg, M. Peifer, Transcription-induced mutational strand bias and its effect on substitution rates in human genes. Mol. Biol. Evol. 26, 131–142 (2009).

48. G. McVicker, P. Green, Genomic signatures of germline gene expression. Genome Res. 20, 1503–1511 (2010).

49. V. Pelechano, L. M. Steinmetz, Gene regulation by antisense transcription. Nat. Rev. Genet. 14, 880–893 (2013).

50. L. J. Core, J. J. Waterfall, J. T. Lis, Nascent RNA sequencing reveals widespread pausing and divergent initiation at human promoters. Science (80-.). 322, 1845–1848 (2008).

51. S. H. C. Duttke et al., Human promoters are intrinsically directional. Mol. Cell. 57, 674–684 (2015).

52. N. J. Proudfoot, Transcriptional termination in mammals: Stopping the RNA polymerase II juggernaut. Science (80-.). 352, aad9926 (2016).

53. H. Menoni et al., The transcription-coupled DNA repair-initiating protein CSB promotes XRCC1 recruitment to oxidative DNA damage. Nucleic Acids Res. 46, 7747–7756 (2018).

54. C. Park, W. Qian, J. Zhang, Genomic evidence for elevated mutation rates in highly expressed genes. EMBO Rep. 13, 1123–1129 (2012).

55. Y. Nédélec et al., Genetic ancestry and natural selection drive population differences in immune responses to pathogens. Cell. 167, 657–669.e21 (2016).

56. H. Quach et al., Genetic adaptation and neandertal admixture shaped the immune system of human populations. Cell. 167, 643–656.e17 (2016).

57. M. Nei, Selectionism and neutralism in molecular evolution. Mol. Biol. Evol. 22, 2318–2342 (2005).

58. Y. Benjamini, Y. Hochberg, Controlling the False Discovery Rate: A Practical and Powerful Approach to Multiple Testing.

59. C. Pál, B. Papp, L. D. Hurst, Highly expressed genes in yeast evolve slowly. Genetics. 158, 927–931 (2001).

60. D. A. Drummond, J. D. Bloom, C. Adami, C. O. Wilke, F. H. Arnold, Why highly expressed proteins evolve slowly. Proc. Natl. Acad. Sci. USA. 102, 14338–14343 (2005).

61. M. Nei, Mutation-driven evolution (books.google.com, 2013).

62. D. E. Barnes, T. Lindahl, Repair and genetic consequences of endogenous DNA base damage in mammalian cells. Annu. Rev. Genet. 38, 445–476 (2004).

63. N. Kim, S. Jinks-Robertson, Abasic sites in the transcribed strand of yeast DNA are removed by transcription-coupled nucleotide excision repair. Mol. Cell. Biol. 30, 3206–3215 (2010).

64. A. Kong et al., Rate of de novo mutations and the importance of father’s age to disease risk. Nature. 488, 471–475 (2012).

65. H. E. Krokan, M. Bjørås, Base excision repair. Cold Spring Harb. Perspect. Biol. 5, a012583 (2013).

66. R. S. Hansen et al., Sequencing newly replicated DNA reveals widespread plasticity in human replication timing. Proc. Natl. Acad. Sci. USA. 107, 139–144 (2010).

67. J. E. Cleaver, Transcription coupled repair deficiency protects against human mutagenesis and carcinogenesis: Personal Reflections on the 50th anniversary of the discovery of xeroderma pigmentosum. DNA Repair (Amst). 58, 21–28 (2017).

68. S. Efroni et al., Global transcription in pluripotent embryonic stem cells. Cell Stem Cell. 2, 437–447 (2008).

69. R. B. Cervantes, J. R. Stringer, C. Shao, J. A. Tischfield, P. J. Stambrook, Embryonic stem cells and somatic cells differ in mutation frequency and type. Proc. Natl. Acad. Sci. USA. 99, 3586–3590 (2002).

70. Y. Hong, R. B. Cervantes, E. Tichy, J. A. Tischfield, P. J. Stambrook, Protecting genomic integrity in somatic cells and embryonic stem cells. Mutat. Res. 614, 48–55 (2007).

## Methods and Supplementary References

71. Valli, H. et al. Fluorescence- and magnetic-activated cell sorting strategies to isolate and enrich human spermatogonial stem cells. Fertil. Steril. 102, 566–580.e7 (2014).

72. Hashimshony, T. et al. CEL-Seq2: sensitive highly-multiplexed single-cell RNA-Seq. Genome Biol. 17, 77 (2016).

73. Dobin, A. et al. STAR: ultrafast universal RNA-seq aligner. Bioinformatics 29, 15–21 (2013).

74. Kodinariya, T. M. & Makwana, P. R. Review on determining number of Cluster in K-Means Clustering. International Journal (2013).

75. Wagner, F., Yan, Y. & Yanai, I. K-nearest neighbor smoothing for high-throughput single-cell RNA-Seq data. BioRxiv (2017). doi:10.1101/217737

76. Baron, M. et al. A Single-Cell Transcriptomic Map of the Human and Mouse Pancreas Reveals Inter- and Intra-cell Population Structure. Cell Syst. 3, 346–360.e4 (2016).

77. Von Kopylow, K. & Spiess, A.-N. Human spermatogonial markers. Stem Cell Res. 25, 300–309 (2017).

78. Chang, Y.-F., Lee-Chang, J. S., Panneerdoss, S., MacLean, J. A. & Rao, M. K. Isolation of Sertoli, Leydig, and spermatogenic cells from the mouse testis. BioTechniques 51, 341–2, 344 (2011).

79. Mali, P. et al. Stage-specific expression of nucleoprotein mRNAs during rat and mouse spermiogenesis. Reprod Fertil Dev 1, 369–382 (1989).

80. Yan, W., Ma, L., Burns, K. H. & Matzuk, M. M. HILS1 is a spermatid-specific linker histone H1-like protein implicated in chromatin remodeling during mammalian spermiogenesis. Proc. Natl. Acad. Sci. USA 100, 10546–10551 (2003).

81. Potter, S. J. & DeFalco, T. Role of the testis interstitial compartment in spermatogonial stem cell function. Reproduction 153, R151–R162 (2017).

82. Ye, L., Li, X., Li, L., Chen, H. & Ge, R.-S. Insights into the Development of the Adult Leydig Cell Lineage from Stem Leydig Cells. Front. Physiol. 8, 430 (2017).

83. Buganim, Y. et al. Direct reprogramming of fibroblasts into embryonic Sertoli-like cells by defined factors. Cell Stem Cell 11, 373–386 (2012).

84. Chen, L.-Y., Willis, W. D. & Eddy, E. M. Targeting the Gdnf Gene in peritubular myoid cells disrupts undifferentiated spermatogonial cell development. Proc. Natl. Acad. Sci. USA 113, 1829–1834 (2016).

85. DeFalco, T. et al. Macrophages contribute to the spermatogonial niche in the adult testis. Cell Rep. 12, 1107–1119 (2015).

86. Rebourcet, D. et al. Sertoli cells modulate testicular vascular network development, structure, and function to influence circulating testosterone concentrations in adult male mice. Endocrinology 157, 2479–2488 (2016).

87. Eden, E., Navon, R., Steinfeld, I., Lipson, D. & Yakhini, Z. GOrilla: a tool for discovery and visualization of enriched GO terms in ranked gene lists. BMC Bioinformatics 10, 48 (2009).

